# The Emergence of Novel Versus Known Three-Dimensional Structures from Random Sequences

**DOI:** 10.64898/2025.12.04.691947

**Authors:** Rose Yang, Hyunjun Yang, Anton Davydenko, Zack Mawaldi, Rian Kormos, Dru Myerscough, Yibing Wu, William F. DeGrado

**Affiliations:** Department of Pharmaceutical Chemistry, Cardiovascular Research Institute, University of California, San Francisco, CA 94158; Department of Biochemistry, Brandeis University, Waltham, MA 02453

## Abstract

It has been hypothesized that while random sequences are unlikely to fold into proteins of the length of globular proteins, repeated random sequences are more likely to adopt stably folded structures, with implications for molecular evolution. We used structure prediction methods to determine the foldability of approximately 120-residue sequences composed of 5-to 60-residue random repeats. With repeats of less than 30-residues, sequences were frequently discovered (1–12%) that fold with high confidence. For less than 60-residue repeats, we frequently observe β-solenoids, similar to those seen in natural proteins. We observe solenoids stabilized by apolar packing as well as ones stabilized by polar interactions with Ca^2+^ in the core of the structure as in natural RTX domains. Helical bundles were observed with high frequency when insertions or deletions (INDELs) were included between blocks of repeating sequences. We also observed a new super-secondary structure consisting of a tightly wound α-helical screw, and experimentally confirmed its stability and structure by CD spectroscopy and X-ray crystallography. Thus, structure predictors can discover structures that are well out of the distribution of the data upon which they were trained. Beyond 40-residue repeat lengths very few sequences were predicted to fold. The small number of structures we observed were representative of well-established major classes of tertiary structures; greater sampling would be needed to discover novel structures from a random distribution. These studies illuminate dark matter regions of protein structure space and support previous predictions that proteins evolved through the assortment of shorter peptide sequences.

**Significance statement:** The availability of powerful and accurate programs for predicting protein three-dimensional structures enables one to ask fundamental questions concerning the origin of folded functional proteins during evolution. We show that 120-residue proteins composed of random sequences repeated in tandem are predicted to be much more likely to fold than fully random proteins. These studies validate previous predictions that proteins evolved through the repetition and assortment of short peptide sequences. Also, some of the predicted structures represent novel conformations, which were confirmed experimentally. These findings advance our understanding of molecular evolution and have implications for design of novel proteins.

## Introduction

Proteins are molecular machines that rely on well-defined tertiary structures to carry out many of their remarkable functions. The molecular mechanisms by which proteins evolved their modern-day sequences and structures have been a puzzle for decades. Early studies focused on the astronomical number of possible sequences^1,2^; for even a 100-residue protein there are 20^100^ possible permutations for a sequence of this length composed of the 20 naturally occurring amino acids. However, more than a single sequence can fold into a single tertiary structure, and there are many possible tertiary structures into which proteins can fold. Moreover, in pioneering work, Margaret Dayhoff identified approximate symmetry in modern-day protein sequences and structures, and suggested that proteins evolved from peptides that assembled into protein-like structures.^3,4^ Gene duplication resulted in repeated sequences in which each individual repeat could evolve independently and asymmetrically under selective pressure for function.^5, 6, 7, 8, 9, 10, 11^ In the resulting structures, the original repeating units are arranged into structures with approximate rotational or dihedral symmetry.^12^ Other repeating sequences are arranged into elongated solenoidal structures with repeat units arrayed in pseudo-translational or screw symmetries. Such repeat proteins are found in enzymes, binding proteins and structural proteins.^9,13,14^

While there have been extensive investigations into design of repeat proteins^15,16^, in each case the design process was biased towards the practical objective of obtaining a predetermined outcome, such as helical repeat proteins^17,18,19^ trefoils^20,21^, or solenoids^22,23,24^. Recent studies have focused on the design of proteins starting from random sequences, using mutations and a guiding function to reach a folded structure.^25,26^ Here, we instead seek to determine the extent to which fully random repeating sequences can fold into native structures without rounds of sequence improvement. What fraction of random sequences are “in-distribution” and hence predicted to fold into tertiary structures already known to occur in the Protein Data Bank (PDB)? We also probe how the probability of folding varies with respect to the length of the sequence repeat, and the presence of insertions and deletions between the repeats. Next, we ask whether there are structures that are very far from the distribution of known structures, but that can nonetheless be identified using programs that are trained on natural proteins. We identify and experimentally characterize a new super-secondary structure.

Historically, designable sequences are defined as ones with an energetically favorable tertiary structure accompanied by a large energy gap between this structure and the next most stable member of the structural ensemble.^27, 28, 29^ Designable structures can accommodate a large number of sequences; the more sequences that can fold into the same structure, the more designable it is defined to be. Here, we determine how repeat length and the presence of INDELs influence the foldability of random sequences.

## Results

### Foldability of random repeat proteins

To estimate foldability, we generated 120-residue proteins composed of repeating units ranging from 5 to 120 amino acids. Two amino acid alphabets were tested: (i) the full set of 20 canonical residues and (ii) a reduced set of 10 “primitive” residues thought to be the ones available early in molecular evolution (G, A, S, T, D, E, I, L, P, V) ^30^. Similar trends were observed for both alphabets up to 40-residue repeats; therefore, unless otherwise noted, we focus on results from the 20-residue alphabet.

For each repeat length, approximately one million structures were predicted using RaptorX^31^, chosen for its speed and accuracy with small proteins.^32^ Models were clustered using a greedy algorithm (see methods) with a 1.0 Å RMSD cutoff, which provided sufficient resolution to separate folds with qualitatively different motifs and φ/ψ angle distributions. Long monomeric helices were frequently predicted but excluded from further analysis. Representative cluster centroid sequences were subsequently re-predicted with AlphaFold3 (AF3)^33^; the RaptorX and AF3 models were generally consistent within 1.0 Å RMSD.

The fraction of sequences predicted to fold (“foldable sequences”) depended strongly on repeat length (**Fig. 1A–F**). We required a value of pLDDT > 90 as a stringent test of foldability. The foldability was 3.6% for pentamers, rose to a maximum of 17.2% for 10-residue repeats, and declined sharply thereafter to 0.01% for proteins composed of two 60-residue repeats and 0.001% ppm for fully random sequences (**Fig. 1G**).

**Figure 1.**
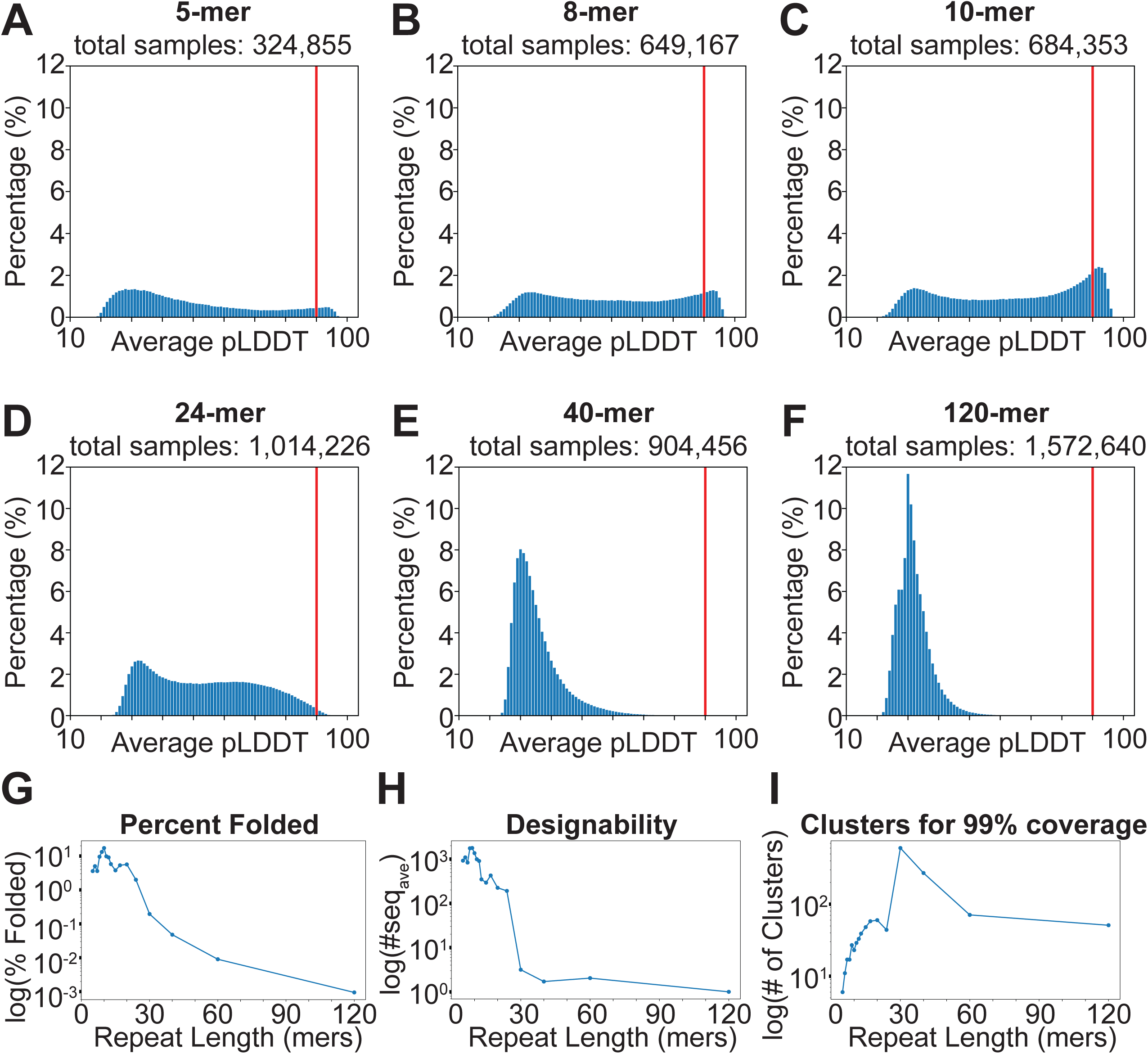
Foldability of random repeat proteins as a function of repeat length. (A-F) Distributions of mean pLDDT for proteins 5, 8, 10, 24, 40 and 120-residue repeats. The red dashed line marks a pLDDT score of 90. (G) Fraction of sequences predicted to fold (pLDDT ≥ 90) per repeat length. (H) Number of structural clusters to achieve 99% coverage of foldable sequences per repeat length. (I) Mean designability of structural clusters (#seq_ave_) per repeat length.

The variation in designability with respect to repeat length (**Fig. 1H**) suggested an interplay between the structural diversity available at each repeat length and the designability of the corresponding folds. To quantify the *structural diversity* available to a given repeat length, we count the number of distinct clusters required to achieve 99% coverage of the total number of foldable sequences for that repeat length. The number of structural clusters increases with the repeat length until a length of 30 residues is reached and declines rapidly thereafter (**Fig. 1I**). We define the *designability* of a given structural cluster as the number of sequences in our sample that are predicted to fold into that structure. We also defined a metric to compare the designability of the full ensemble of structures associated with each repeat length (#seq_ave_, **Fig. 1H**) as the mean number of sequences in each cluster for that repeat length. For short repeat lengths of less than 13-residues, a small number of highly designable structures dominated the foldable space, resulting in high values of #seq_ave_.

Together, these data show that there are relatively few designable conformations for structures with short repeat lengths (**Fig. 1H**), but this number increases as the number of mainchain torsional degrees of freedom increases. However, at repeat lengths longer than 30 residues, foldable sequences occur with such low frequencies that we observe far fewer within our sample size of one million sequences/repeat length.

### Sequence and structural features of highly designable structural repeats

*Pentameric repeats.* For pentameric repeats, the sequence space is small enough (approximately 20^5^ / 5 = 640,000 unique sequences) to be exhaustively sampled (**Fig. 2**). Of these, 3.6% were predicted to fold into one of five β-solenoid–like structures that together account for ∼98% of foldable sequences (pLDDT > 90, **Fig. 2A**). The two dominant clusters (cluster_0 and cluster_1, **Fig. 2B, D**) are structurally very similar, and together comprised ∼89% of all models. They both form right-handed, square β-solenoids (4.00 ± 0.05 repeats/turn and a pitch of 4.6 Å/turn) formed by repeating β-strands connected by sharp turns. The trajectory of the chain at the vertices of the squares resembles that of a classic turn (**Fig. 2F, G**): a mix of type I and type II β-turn conformations are found in cluster_0 and predominantly type I β-turns in cluster_1.

**Figure 2.**
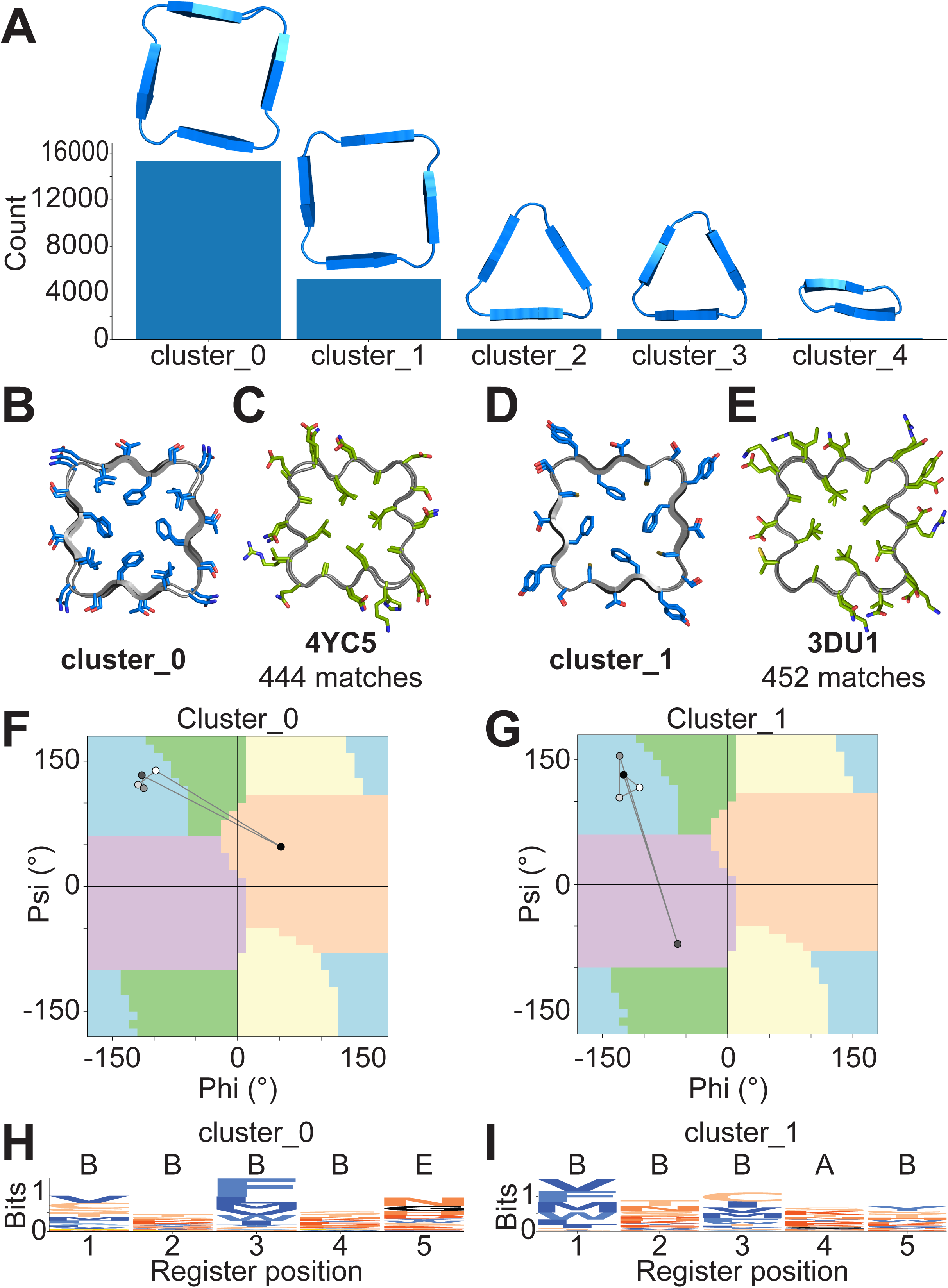
Features of 5mer repeat proteins. (A) Histogram depicting frequency and structure of each cluster. (B, D) Cross-sectional models of the two dominant pentameric clusters. (F, G) Plots of φ/ψ angles of centroids highlighting β-strand geometries (positions 1–3) and turn conformations (positions 4–5). (C, E) Closely matching natural β-solenoids from the PDB. (H, I) Sequence logos for the clusters.

Database searches confirmed that the centroid structures closely matched known β-solenoids in natural proteins (**Fig. 2C, E**). The geometry of the 5-mer clusters 0 and 1 were observed 444 and 452 times, respectively, indicating that these motifs occur frequently in nature. The sequence logos for both natural and random pentameric repeats showed similar hydrophobic enrichment at positions 1 and 3, consistent with packing of side chains into the solenoid core (**Fig 2H, I**). ^14,23,34,35,36^ Finally, predictions restricted to the 10 “primitive” amino acids yielded the same four structures, although triangular solenoids were relatively more common than square structures in this reduced alphabet (**Fig. S1**).

*Hexameric repeats.* For repeats longer than 5 residues, exhaustive enumeration of sequence space was no longer feasible with our computing resources, so we analyzed approximately one million random sequences. 5.0% of these sequences were folded into one of 10 different solenoidal topologies (**Fig. 3**). The three largest clusters are triangular and account for **83.5**% of the random sequences analyzed; square and oval β-solenoids were much less common (**Fig 3A**). The two most frequent clusters are right-handed solenoids with a triangular cross-section (3.0 residues/turn) and an axial rise of approximately 4.5 Å/turn (**Fig. 3B, D**). The β-strands are one residue longer in the 6-mers than in the above 5-mers. The corners are formed by type II and type I turns in clusters 0 and 1, respectively, which were enriched in glycine (**Fig. 3F, G**). The structures were qualitatively similar to known structures of repeating hexamers and database searches confirmed structural similarity to natural proteins, returning 435 and 75 hits for clusters 0 and 1, respectively (**Fig. 3C, E**). In both natural and the randomly generated sequences, hydrophobic residues are strongly enriched at positions 1 and 3 of the β-strand segments, packing into a central apolar core (**Fig. 3H, I**).

**Figure 3.**
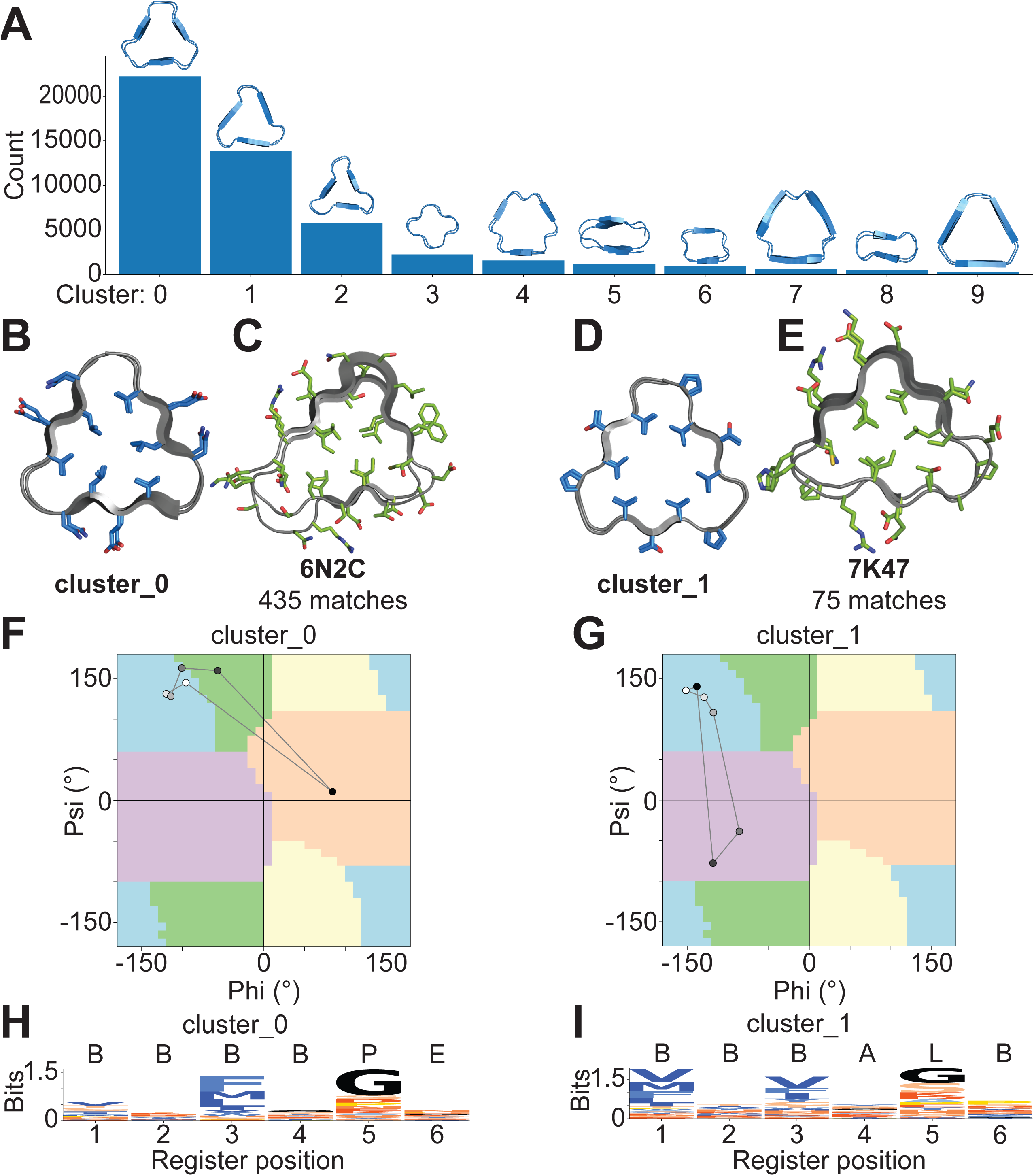
Features of 6mer repeat proteins. (A) Histogram depicting frequency and structure of each cluster. (B, D) Cross-sectional structures of the two dominant pentameric clusters. (F, G) Centroid Ramachandran plots highlighting conserved β-strand φ/ψ geometries (positions 1–3). (C, E) Closely matching natural β-solenoids from the PDB. (H, I) Sequence logos for the clusters.

*7–10 residue repeats.* Repeat lengths in the 7–10 residue range marked a transition from compact uniform, β-solenoids toward more diverse, elongated and structurally adaptive solenoids. Heptameric and nonameric repeats produced relatively high foldability fractions (3.6% and 13.2%, respectively). Compared to the 5- and 6-mers, their folds were more numerous and varied: In 7-mers, a triangular arrangement with three repeats per turn was occasionally observed, but sidechain packing was looser than in hexameric solenoids. In both 7- and 9-mers, β-arcade geometries were common, in which two repeat units formed paired β-arches.^14,37,38^ The cores were stabilized by apolar sidechains, though in some clusters internal Asn and Gln residues contributed hydrogen-bonded zippers.^39,40,41^

Even-numbered repeats tended to form symmetrical rectangular assemblies whose long edges are regular β-strands. Eight-residue repeats (**Fig 4A, B**) produced two major rectangular clusters (∼45% of models) with tight arches forming the short edges. Consistent with their rectangular symmetry, they have 2.0 turns/repeat and an axial rise of 4.8 Å. Ten-residue repeats (**Fig 5A, B**) tended to have a rectangular cross-sectional β-solenoid motif (2.0 repeats/turn), which is elongated or widened relative to the 8-mer.

**Figure 4.**
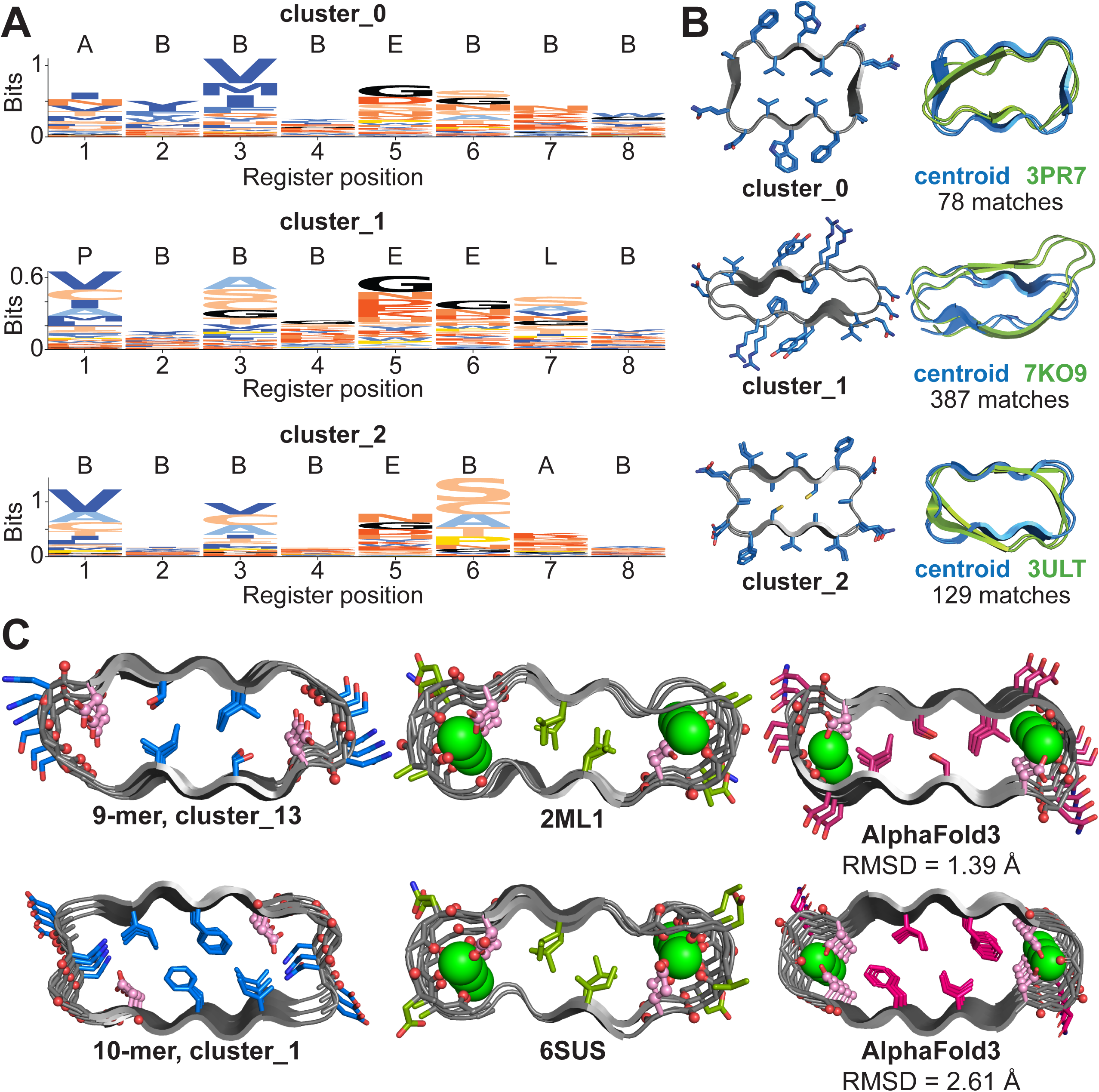
Features of 8mer repeat proteins. (A) Sequence logos and secondary structure assignments for the three largest clusters. (B) Axial views of structures and comparisons with natural proteins. (C). Members of 9mer and 10mer repeat clusters that resemble natural RTX calcium-binding motifs, with AlphaFold3 predictions likewise positioning Ca²⁺ at acidic residues.

**Figure 5.**
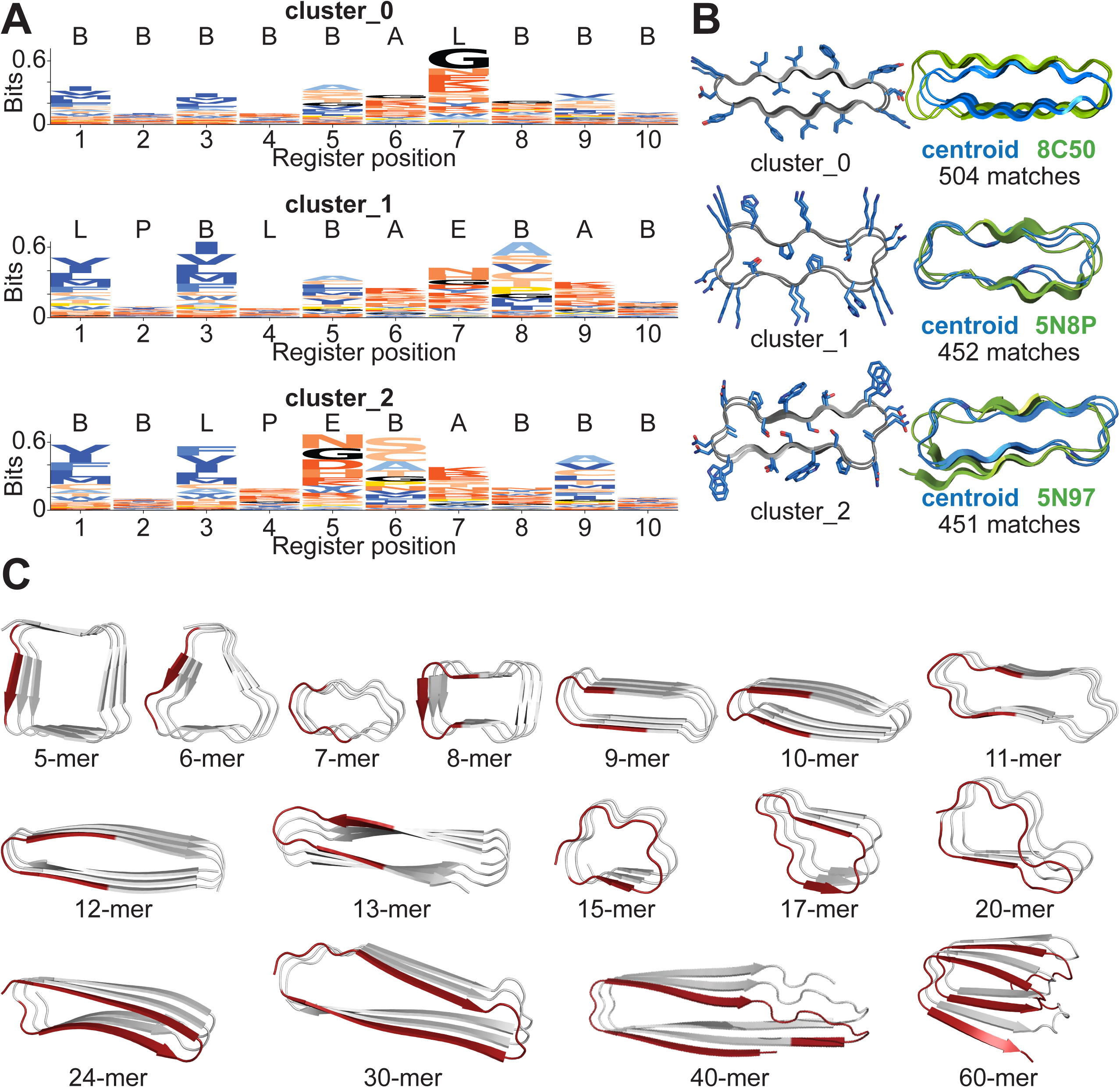
Features of 10mer repeat proteins. (A, B) Sequence logos and cross-sectional structures for the three major clusters. (C) Centroid structures for repeats ranging 5 – 60 residues. As repeat length increases, the number of repeats per turn (red) decreases.

While the cores of the 9-mers and 10-mers were generally well packed with hydrophobic (Ala, Val, Ile, Leu, Met, Phe) or Ser/Thr, we also observed Asn or Gln residues with their sidechains forming extensive sidechain-sidechain and sidechain-mainchain, as seen in natural amyloids and solenoids.^39,40,41^ We also were initially surprised to see that members of 9-mer and 10-mer clusters had Asp residues pointing inward proximal to several mainchain carbonyl groups. AlphaFold is known to predict metal-binding sites, even without the metal ions included, and the oxygen-rich environment appeared ideal for binding Ca(II) ions. Indeed, a search of the PDB identified repeating RTX Ca(II)-binding motifs and enzymes (e.g., 1RU4) with closely related structures in 7788 proteins. RTX motifs play an important role in bacterial secretion. Thus, our workflow identifies metal ion-binding motifs as well as structural motifs.

Together, these findings highlight a progression in which the number of repeats required to complete a turn of the solenoid decreases with increasing repeat length: 5-mer repeats form squares with four repeats/turn, 6-mers form triangles with three repeats/turn, 8-mers to 12-mers form increasingly variable shapes with two repeats/turn; 15–20-mers have a single repeat/turn, and 40 residue repeats have 0.5 repeats/turn (**Fig. 5C**). As a result, the number of residues per turn of the solenoid stays roughly constant between 15 and 30 residues. This behavior appears to be dictated by the need to maintain efficient packing of interior sidechains. The emergence of greater geometric diversity at longer repeat lengths reflects increased conformational freedom as the number of backbone torsional angles increases. We also note that although both right-handed and left-handed β-solenoids are seen in known structures, with 70% frequency representing right-handed structures^14^, we found almost exclusively right-handed solenoids in our models. This likely reflects the fact that the great majority of monomeric solenoids are right-handed, while solenoids that associate to higher order structure are often left-handed. Additionally, it has been noted that AlphaFold2 tends to predict right-rather than left-handed solenoids for self-associating β-solenoids known to be left-handed from independent structural studies.

### Discovery of a novel structural unit comprising a tightly wrapped α-helical screw

Interestingly, we also observed a novel super-secondary structure, which we define as an α-helical screw, in which a series of interconnected helices are tightly wrapped around a central axis. Helical screws were observed most frequently in 8-mer repeats (133 occurrences); they were also present in 7-mer (30 occurrences) and to a lesser extent in other repeat lengths (**Fig. 6A, B**). We observe variations on this basic motif, which differ in how tightly the superhelix is wound (**Fig. 6D**). We identify and name a given screw conformation based on its repeat length and the number of repeats per turn (e.g., an 8-mer_2.5_ would have 2.5 8-mer units per superhelical turn). We observe a range of 2.5 to 4.0 units per turn of the helical screw, and all were found to wind in a clockwise manner. A TM-align^42^ search of the PDB revealed only a single example of an 8-mer helical screw (**Fig. 6C**), and no hits were found for other helical screw conformations. Thus, helical screws represent a “dark-matter” portion of protein conformational space.^43,44^

**Figure 6.**
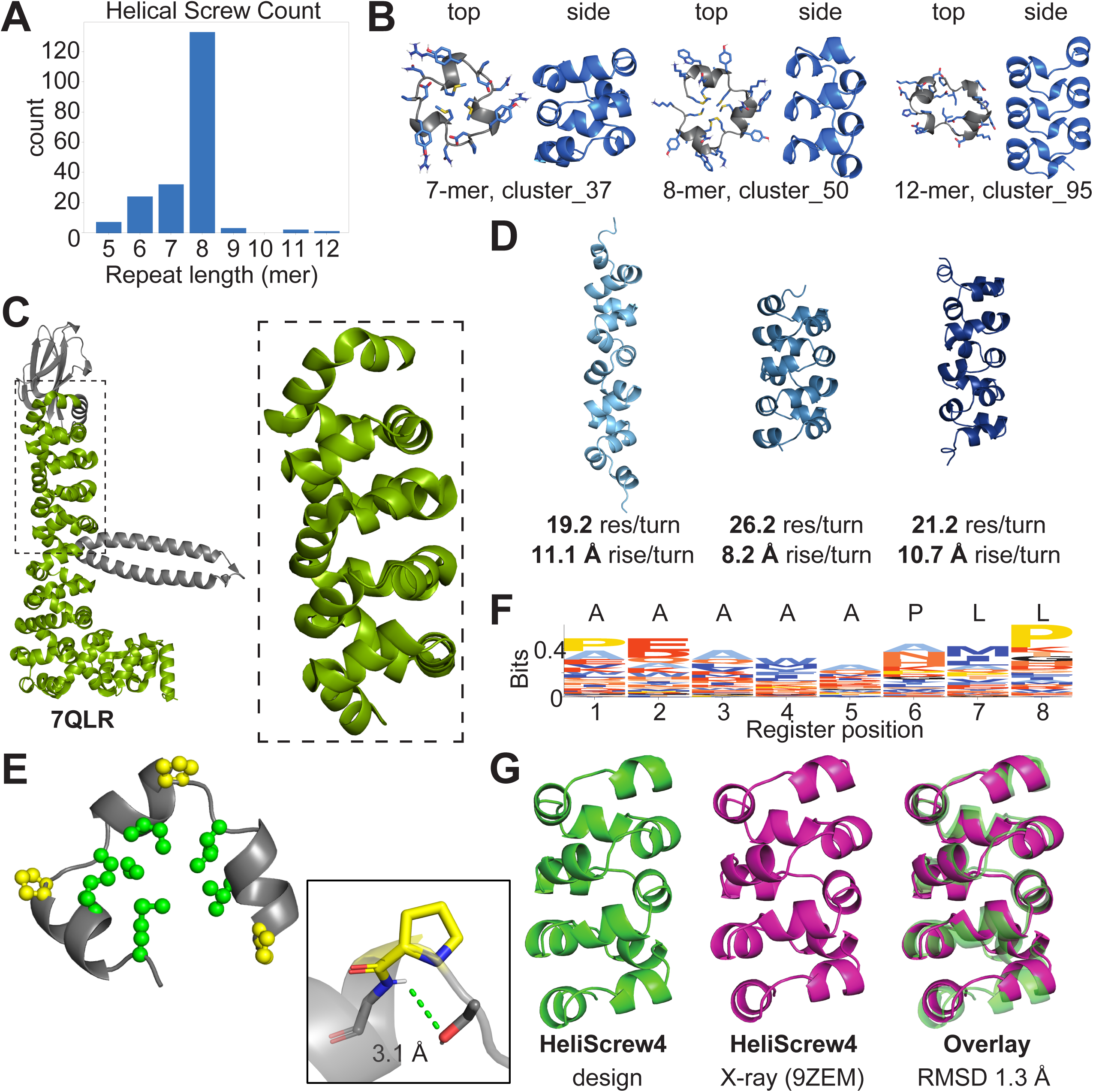
(A) Frequency of occurrence of α-helical screws as a function of repeat length. (B) Centroid structures from the largest clusters for 7mer, 8mer, and 12mer repeats. (C) Structure of the natural protein 7QLR containing a similar helical screw. (D) Examples of three distinct α-helical screw geometries observed for 8mer repeats. (E) Cross-sectional structure of a member of the largest 8-mer cluster. Hydrophobic core-packing residues (green) and a helix-initiating Pro (yellow) are shown. (F) Sequence logo representation of the 8mer cluster with APBLE annotation^63^. (G) Comparison of experimentally determined structure HeliScrew4 (magenta) with the design model HeliScrew7 (green), showing close agreement (b.b. r.m.s.d. = 1.38 Å).

The 8-residue helical screws are all stabilized by hydrophobic clustering of apolar sidechains in the core of the superhelical screw, as seen in the predicted structures and sequence logos of the motif (**Fig. 6E, F**). Moreover, Pro occurs frequently at the N1 position of the helix, and short polar residues Ser, Thr, Asp, Asn occupy the N-cap positions where they form hydrogen bonds to the amides exposed at the N-terminus of the helices.^45^

Given the novelty of this structural type, we sought to experimentally characterize an 8-mer helical screw, which we term HeliScrew. To reduce sequence redundancy and ensure a water-soluble exterior, we used Chroma^46^ to design six helical screw sequences (HeliScrew1–6) derived from the original cluster centroid structure (HeliScrew7). All seven proteins expressed robustly in *E. coli* and were soluble in aqueous buffer at pH 7.5 by SDS-PAGE and SEC FPLC (**Fig. S22–S25**). All constructs eluted as apparent monomers except HeliScrew6, which showed a shifted higher–molecular weight peak by SEC–FPLC, indicative of higher-order assembly.

All seven proteins were subjected to X-ray crystallization screening. HeliScrew4 crystallized in 1.8 M ammonium citrate tribasic, pH 7.0, yielding rod-shaped crystals suitable for X-ray diffraction. Data were collected at Argonne National Laboratory (APS beamline 24-ID-E). The structure was solved by molecular replacement and refined in space group P12_1_1 to 2.20 Å resolution. The asymmetric unit contains two crystallographically independent HeliScrew4 molecules, one of which is shown in **Fig. 6G**. Comparison of the experimentally determined HeliScrew4 structure (magenta) with the Chroma design model (green) shows close agreement (Cα backbone r.m.s.d. = 1.05 Å; **Fig. 6G**). Comparison of the HeliScrew4 crystal structure with the cluster centroid HeliScrew7 also shows close agreement (Cα backbone r.m.s.d. = 1.38 Å). Circular dichroism spectroscopy of HeliScrew4 revealed a typical α-helical signature with negative minima at 208 and 222 nm (**Fig. S26**), suggesting that the protein adopts the designed α-helical structure in solution.

The high degree of “expressibility” and water solubility of these helical screw sequences as well as the excellent correspondence between the predicted and the crystallographically observed α-helical screw structures shows that they are indeed highly designable examples of structures from the “dark matter” region of conformational space.

### Globular folds emerge in 60-mer and non-repeating structures

Beyond a repeat length of 40 residues, there is an abrupt change from repeating solenoidal structures to globular structures –all with folds that are well represented in the PDB. For 60-mers we observed a mixture of all-α folds, all-β, and α/β structures (**Fig. 7A**). All represented common topologies, and we found that their tertiary structures were covered to an extent of 80% or greater using two CATH domains^47^ using CATHcover (**Fig. 7B**). No sequences were predicted to fold with a pLDDT > 90 for the fully random 120-mer, although 11 structures were found to have values of pLDDT > 80. Their structures included 3 helical bundles, 1 all-β structure and 7 α/β structures (**Fig. S27**), and all were covered at 80.0% percent or more by 2 or 3 CATH domains.

**Figure 7.**
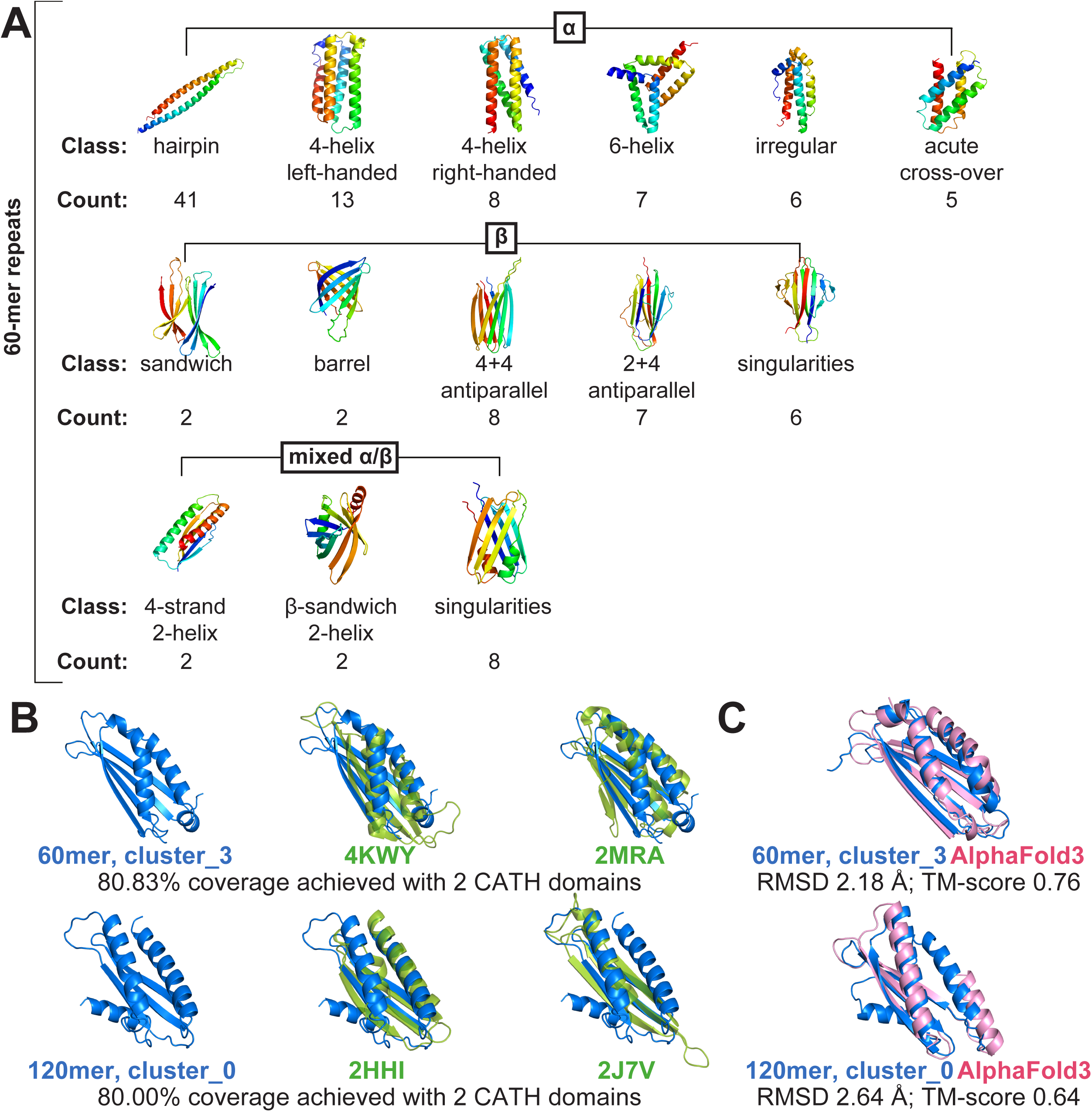
(A) 60mer repeat proteins with α, β, and mixed α/β architectures, and the number of structures observed in each cluster. (B) Comparisons between designed repeat proteins and their closest natural CATH domains. Centroids for 60mer_cluster_3 and 120mer_cluster_0 align closely with natural folds, achieving ∼80% coverage with two CATH domains. (C) Overlays with AF3 predictions.

### Insertions and deletions encourage formation of a-helical bundles

We observed helical structures with much lower frequency than for β-solenoids. Helical hairpins, with two helices running anti-parallel to one another were seen 393 times for 7-mer repeats (1.1% of the folded sequences) and 286 times (0.3%) for 11-mer repeats. Three- and four-helix bundles were even more rare, being observed only in the 40-mer repeats (6 times, 1.4%), and the 30-residue repeats (25 times, 1.3%), respectively.

We were initially surprised by the relative paucity of conventional helical bundles and coiled coils in our random proteins, given that they occur widely in natural proteins. Instead, sequences that might have been expected to form coiled coils were predicted to form extended, monomeric α-helices. However, when we assessed the *intermolecular* folding of sequences composed of random 7-mer repeats as non-covalent homodimers, we found 15.4% of the sequences had a value of pLDDT > 90. The corresponding values for 3-, 4-, 5- and 6-stranded structures were 3.5%, 6.8%, 1.2%, and 0.3%, respectively.

These calculations suggested that intramolecularly folded helical bundles might be formed with high frequency only if one allowed occasional breaks between blocks of heptad repeats to allow the protein helices to fold back on one another. To test this idea, we systematically introduced INDELs, consisting of either 1- or 2-residue deletions or 1 to 4-residue Gly insertions between blocks of four repeated random heptads (to create app. 28-residue helices). This pattern was repeated four times to create potential 4-helix bundles. The presence of deletions indeed increased the fraction of sequences that folded with high confidence by 20-fold, from 0.002% to approximately 0.04%. More dramatic, 300 to 500-fold increases were seen with 1 to 4-residue Gly insertions, reaching 0.93% for a two-residue insertion (**Fig. 8A**). The large majority (39.7%) of the predicted structures represented classical left or right-turning helical bundles (**Fig. 8B, C**). Similar results were seen for sequences intended to form 5- or 6-helix bundles, respectively, although they were found to fold with lower frequency, as expected from the fact that they occur less frequently than 2, 3, or 4-helix bundles in nature (**Fig. S34, S35**).

**Figure 8.**
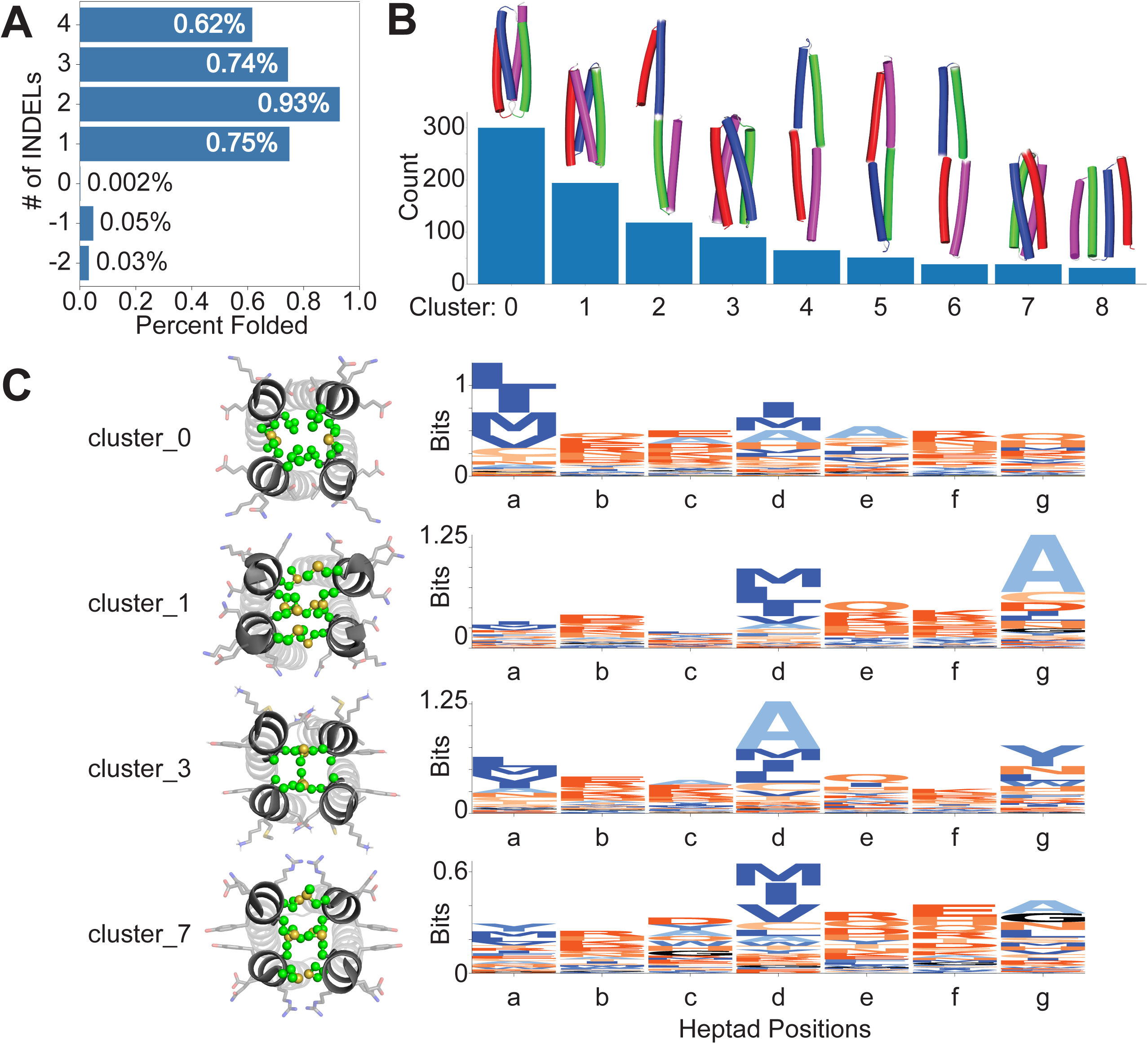
Effect of INDELs on the monomolecular folding of sequences with blocks of four, 7-residue repeats. (A) The percentage of sequences that folded into helical bundles as a function of the indel type. (B) Frequency and structure of each cluster. (C) Sequence logos for the largest cluster with 2-Gly insertions. Clusters 0, 1, 3 and 7 are left- and right-turning 4-helix bundles, while the remaining structures contain helical hairpin units. Hydrophobic residues at positions 1 and 4 project tend to project towards the interior of each 4-helix bundle.

## Conclusions

Our results show that the inclusion of repeats enriches the fraction of otherwise random sequences that can adopt folded conformations. With repeat lengths of 5 to 20 residues, over 1% of the sequences are predicted to fold, mostly into repeating solenoids or β-hairpins, although we also observe helical bundles and tightly wound helical screws that represent a new family of super-secondary structures. With the inclusion of INDELs between blocks of repeats, we also observe the formation of helical bundles at a frequency of 0.5 to 1%.

On the other hand, beyond a repeat length of 30 residues, the frequency of observing predicted folded structures decreased sharply, reaching 0.001% for sequences lacking repeats. Also, these structures were all globular. Evidently, without the imposition of a sequence repeat, the probability of adopting a repeating conformational form becomes very low, and only more asymmetric arrangements typical of the domains seen in globular proteins are observed. The resulting structures are representative of ones seen in globular proteins, and include the α-helical, α/β, and all-β classes of proteins.

These observations answer our first questions concerning how frequently random sequences are likely to fold into structures similar to those found in nature. However, this does not necessarily guarantee that all predicted proteins would fold as monomeric species in aqueous solution. In nature, solenoids and repeating β-hairpins are capped on at least one end to prevent end-to-end polymerization. In the absence of such repeat-breaking features we expect that proteins discovered from precisely repeating sequences would be prone to forming polymeric structures similar to amyloid fibrils. Indeed, short, amyloid-forming peptides have been hypothesized to have been important players in the molecular evolution of proteins.^48,49,50^ They are known to have catalytic functions^41,51,52,53^ and they can form membrane-like tubes and spheres as well as phase-segregated states, ^54, 55, 56, 57^. A solenoid combines many copies of monomeric amyloidogenic peptides into a single chain, allowing each monomer to mutate independently to evolve function. Indeed, solenoids form the frameworks of many enzymes across multiple catalytic classes, and they serve as structural domains in multi-domain proteins, anti-freeze proteins, cell-surface proteins, and viruses.^35^

We also noted that up to half of the predicted structures do not appear to have sufficiently polar exteriors to maintain water-solubility. Some of the helical bundle proteins had fully hydrophobic exteriors, which would make them good candidates for insertion into membranes. On the other hand, we are unaware of any transmembrane solenoids and so we expect that a large fraction of the predicted structures would be insoluble if expressed or chemically synthesized. Thus, the predictions shown here should be considered upper limits of the fraction of proteins that would be folded and soluble in membranes or aqueous solution.

We, like previous investigators ^58^, were also initially concerned about the presence of polar residues in the cores of the predicted structures of solenoids. While these features were previously considered to be indicative of false predictions, closer inspection indicated that the polar residues could impart function. For example, one prominent feature was a series of interior-facing Asp residues, that comprise the Ca(II)-binding repeats in natural RTX motifs^54,59,60^ Other polar residues in the interior include Asn residues, which are common in zippers that stabilize parallel β-sheet structures and buried salt bridges, which are common in both water-soluble solenoids and amyloids. We also observe clusters of His, Glu/Asp and Cys residues that could bind transition metals known to catalyze hydrolytic and oxidative reactions. Indeed, metallo-amyloids that contain His-Xxx-His and related motifs form highly active esterases, carbonic anhydrases, and oxidases.^41,53,61^

Our next question was whether structure prediction programs would be able to extrapolate from the distribution found in the PDB to identify new folds. Previously, folding predictors have been used during the design of proteins that have structures with arrangements of secondary structures not frequently or ever seen in the PDB.^43,44^ Here, we discovered an entirely new family of tightly wound helical screws. We observed different conformers for different helical screw repeat lengths, forming distinct sub-families with characteristic superhelical parameters. Only a single member of one of the helical screw family had previously been seen in nature, which prompted us to express and determine the structure of a *de novo* helical screw. The structure is in close (< 1.38 Å backbone RMSD; < 1.58 Å all heavy atom RMSD) agreement with the predicted structure, indicating that the motif is designable. Why then, do we not see it frequently in natural proteins? The sequence requirements required to adopt this structure appear to be restrictive: the core is tightly packed with sterically restricted apolar side chains, and the helix is generally initiated by strong helix-promoting sequence, including a helix-initiating Pro near the N-terminus. It is possible that the sequence requirements for folding are so restrictive that there is little ability for variation in the sequence to generate function.

Our remaining question is whether there are other stably folded structures that are not predicted by current prediction methods. Any structure that is highly extended would have been removed by our radius of gyration filter, which we found necessary to eliminate very long helices or extended structures. Beyond this limitation, the question hinges on whether the set of structural fragments and the interactions required for assembling them into higher order structure represent a covered set. Grigoryan and coworkers have argued that a complete set of fragments and instructions to assemble them is largely covered in the PDB.^29,44, 62^ Indeed, at the level of sequentially contiguous backbone structures, it is likely that all energetically viable sequentially local structures (such as α-helices, hairpins, omega loops, polyproline II helices…), and the way that these fragments can interact is limited by simple physical forces that are captured in folding predictors. Thus, it is likely that—within our sampling size limit—we have identified most if not all possible designable folds up to lengths of approximately 30 residues. Beyond this length, our sampling is too low to identify rare folds, and the few folds we find are well precedented in the PDB. Given our discovery of a completely novel super-secondary structure in the regime where we do have good sampling, it is reasonable to expect that there might be other folds that would be discovered in a larger sample. Such hypothetical structures would not be part of the modern repertoire for a variety of reasons: 1) they might have not been sampled during evolution, 2) they might be refractory to experimental methods of structure determination, 3) they might have been sampled during evolution but failed to confer a functional advantage, or 4) they might possess an inherent liability, such as toxicity or a tendency to misfolding. New tertiary structures might, however, confer advantages for applications or environments not currently found in nature.

## Methods

### RaptorX random repeat generation

Each n-residue repeat was repeated in tandem to produce a 120-residue chain, with truncation or extension applied when the 120/n was not an integer. Repeat lengths of 5 - 13, 15, 17, 20, 24, 30, 40, 60, and 120 residues were analyzed. All sequences were submitted to RaptorX on UCSF’s Wynton high-performance compute cluster, and models with mean pLDDT ≥ 90 were retained. For the primitive-alphabet analysis approximately 250,000 sequences per repeat length were generated.

### Clustering method

A single model was selected at random as the initial seed, and all other structures were compared to it using TM-align (v2020). For every comparison, the full 120-amino acid sequence was used. Models with RMSD ≤ 1.2 Å and TM-score ≥ 0.7 relative to the seed were assigned to the same cluster. A new unassigned model was then chosen as the next seed until all structures were classified. Cluster sequences and their predicted structures are available as **supplementary data**.

### Structure analysis

Aligned sequences were reconstructed from TM-align output logs. Consensus sequences were generated by applying a column-wise majority rule to the aligned sequences. Cluster centroid structures were analyzed using Bio.PDB (Biopython v1.83) to calculate φ and ψ dihedral angles across repeat segments. Torsion angles were projected onto APBLE Ramachandran map assignments to classify local backbone preferences, including α-helical, β-sheet, left-handed helical, and extended regions.^63^ Position frequency matrices were converted to sequence logos, with amino acids colored according to a fixed scheme described in supplementary methods. For helical repeat motifs, residues were annotated using heptad notation (A–G).

### Protein design

Initial structural models for protein design were derived from the cluster centroid of helical screws from the largest 8-mer cluster. Each starting backbone (74 total centroids) was first perturbed by adding noise (t = 0.5) using partial diffusion module in Chroma^46^, followed by denoising and subsequent sequence design steps. For each seed, three designed sequences were generated, yielding 296 sequences in total (74 original + 222 designed). All sequences were evaluated by ESMFold^64^ to obtain predicted structures. The designed models were chosen from the top 25^th^ percentile of ESMFold pLDDT scores and top 25^th^ percentile of backbone agreement between design and prediction (Cα RMSD). Only models passing both thresholds were chosen, preserving a pool of 74 top-scoring sequences per iteration. This iterative diffusion–design–prediction loop was repeated for 20 cycles, yielding 74 structures that converged toward high-confidence models with high pLDDT and low RMSD. The seven top ranking designs (HeliScrew1–6) and the original cluster centroid (HeliScrew7) were selected for expression and experimental characterization. Sequences for all designed models are provided in the supplementary data (**Fig. S22)**.

### Protein expression and purification

Genes encoding the seven design proteins and the original cluster centroid protein (Twist Bioscience) were cloned into a pET-16b expression vector (Amp marker; BamH1 and Not1) containing an N-terminal His-tag-SUMO fusion tag and a cleavage site for human SENP2 protease. Gene sequences are provided in supplementary data (**Fig. S23**). Constructs were transformed into BL21(DE3) *E coli* cells, which were streaked onto LB agar plates. A single colony from each transformation was inoculated into LB medium, and protein expression was induced with 0.5 mM IPTG at an OD of ∼0.5. Cultures were harvested after 4 h of induction at 37 °C. Cell pellets were collected by centrifugation and resuspended in lysis buffer (50 mM NaPhos, 150 mM NaCl, pH 7.5), followed by sonication. Lysates were clarified by centrifugation and loaded onto nickel-nitrilotriacetic acid (Ni-NTA) agarose bead affinity columns for immobilized metal affinity chromatography (IMAC). Bound proteins were eluted using 250 mM imidazole containing lysis buffer. Eluted proteins were dialyzed overnight to remove imidazole and subsequently cleaved with recombinant human SENP2 protease to remove the His-SUMO tag, leaving a single additional serine at the N-terminus. Cleaved proteins were separated by a second round of IMAC purification. Final purification was performed using size-exclusion chromatography (SEC) FPLC on a Superdex 200 Increase 10/300 GL column equilibrated in 50 mM sodium phosphate, 150 mM NaCl, pH 7.5. Peak fractions corresponding to monomeric species were pooled and concentrated for downstream biophysical characterization, including CD spectroscopy and X-ray crystallography.

### Protein crystallization

Crystallization trials were set up for all seven helical screw proteins, testing 384 conditions (Hampton screen Index, Crystal, PEG, JCSG). Heliscrew4 formed in 1.8 M ammonium citrate, pH 7.0; crystallization by hanging drops gave X-ray diffraction-grade crystals. Diffraction data were collected at the Advanced Photon Source, Beamline 24-ID-E (NE-CAT; 12.66 keV; 0.2 ° oscillation over 360° (**Table S2**, pdb 9ZEM). The structure of the helical screw was solved by molecular replacement using the designed model in Phenix.phaser ^65^ . Iterative model building and refinement was carried out in Phenix.autobuild ^66^, Coot ^67^, and Phenix.refine^68^.

### Circular Dichroism (CD)

Far UV CD spectra were recorded using a Jasco J-810 instrument for Heliscrew4 in H_2_O) in a 1 mm pathlength cell. Spectra were collected in 0.5 nm increments from 260–190 nm at a rate of 50 nm/min; 3 scans were averaged.

## Acknowledgements

W.F.D. acknowledges support from the National Science Foundation (NSF, CHE-2528384 and MCB-2306190) and the National Institutes of Health (R35GM122603). H.Y. acknowledges support from the National Institutes of Health (R00AG084926). X-ray diffraction studies were performed under APS beam time award (DOI: https://doi.org/10.46936/APS-191095/60014853) from the Advanced Photon Source, a U.S. Department of Energy (DOE) Office of Science user facility operated for the DOE Office of Science by Argonne National Laboratory under Contract No. DE-AC02-06CH11357. H.Y. thanks Dr. Kay Perry for beamline support. Z.M. acknowledges support from the NIH (NIH T32GM149436). D.M. acknowledges support from the NSF Graduate Research Fellowships Program (GRFP).

## Data availability

All sequences and computed structures for the proteins are available here. (Dataset DOI: 10.5061/dryad.vmcvdnd4m). Statistics concerning the size of clusters and pdb codes and chain/sequence identifiers are provided in the **supplementary data.**

## Code availability

Code to cluster the predicted structures are available here. (https://github.com/degrado-lab/repeating-sequences.git)

## Author contributions

R.Y., H.Y., Y.W., and W.F.D designed the research; R.Y., H.Y., R.K. A.D. Z.M., D.M. and Y.W. generated the structures. R.Y., H.Y., Y.W., and W.F.D analyzed the data; and R.Y., H.Y., Y.W., and W.F.D wrote the paper. Y.W. and W.F.D. supervised the study. All authors read and approved the manuscript.

## Competing interests

The authors declare no competing interests.

## Notes

https://github.com/degrado-lab/repeating-sequences

https://datadryad.org/dataset/doi:10.5061/dryad.vmcvdnd4m

### Competing Interest Statement

The authors have declared no competing interest.

### Summary of Updates

Authorship list and corresponding authors have been updated.

