## Supplemental Information for "The Emergence of Novel Versus Known Three-Dimensional Structures from Random Sequences"

**This PDF file includes:**

| **Generation and Structure Prediction of Random Repeat Sequences** | S3 |
| --- | --- |
| **Primitive amino acid structure generation** | S5 |
| **Cluster Analysis Pipeline** | S6 |
| **Structural Database Searches** | S9 |
| **Fig. S1.** Conformational clustering and sequence features of pentameric repeat proteins generated from “primitive” amino acid set. | S10 |
| **Fig. S2.** Conformational clustering and sequence features of hexameric repeat proteins generated from “primitive” amino acid set. | S11 |
| **Fig. S3. Conformational clustering and sequence features of designed 7-mer repeat proteins generated from the “primitive” amino acid set.** | S12 |
| **Fig. S4. Conformational clustering and sequence features of designed 7-mer repeat proteins generated from the full 20 canonical amino acid set.** | S13 |
| **Fig. S5. Conformational clustering and sequence features of designed 8-mer repeat proteins (early 10 set).** | S14 |
| **Fig. S6. Conformational clustering and sequence features of designed 8-mer repeat proteins generated from the full amino acid set.** | S15 |
| **Fig. S7. Conformational clustering and sequence features of designed 9-mer repeat proteins (early 10 set).** | S16 |
| **Fig. S8. Conformational clustering and sequence features of designed 9-mer repeat proteins generated from the full amino acid set.** | S17 |
| **Fig. S9. Conformational clustering and sequence features of designed 10-mer repeat proteins (early 10 set).** | S18 |
| **Fig. S10. Conformational clustering and sequence features of designed 10-mer repeat proteins generated from the full amino acid set.** | S19 |
| **Fig. S11. Conformational clustering and sequence features of designed 11-mer repeat proteins (early 10 set).** | S20 |
| **Fig. S12. Conformational clustering and sequence features of designed 11-mer repeat proteins generated from the full amino acid set.** | S21 |
| **Fig. S13. Conformational clustering and sequence features of designed 12-mer repeat proteins (early 10 set).** | S22 |
| **Fig. S14. Conformational clustering and sequence features of designed 12-mer repeat proteins generated from the full amino acid set.** | S23 |
| **Fig. S15. Conformational clustering and sequence features of designed 13-mer repeat proteins (early 10 set).** | S24 |
| **Fig. S16. Conformational clustering and sequence features of designed 13-mer repeat proteins generated from the full amino acid set.** | S25 |
| **Fig. S17. Conformational clustering and sequence features of designed 15-mer repeat proteins generated from the full amino acid set.** | S26 |
| **Fig. S18. Conformational clustering and sequence features of designed 17-mer repeat proteins generated from the full amino acid set.** | S27 |
| **Fig. S19. Conformational clustering and sequence features of designed 20-mer repeat proteins generated from the full amino acid set.** | S28 |
| **Fig. S20. Conformational clustering and sequence features of designed 24-mer repeat proteins generated from the full amino acid set.** | S29 |
| **Fig. S21. Conformational clustering and sequence features of designed 30-mer repeat proteins generated from the full amino acid set.** | S30 |
| **Fig. S22. Amino acid sequences of HeliScrews (1–7).** | S31 |
| **Fig. S23. Codon optimized DNA sequences encoding HeliScrew1–7 used for recombinant expression in *E. coli*.** | S32 |
| **Fig. S24. SDS-PAGE analysis of purified HeliScrew1–7.** | S33 |
| **Fig. S25. Size-exclusion chromatography (SEC) traces of HeliScrew1–7.** | S34 |
| **Fig. S26. Far-UV circular dichroism spectrum of HeliScrew4.** | S35 |
| **Fig. S27.** Eleven designed 120-mer proteins predicted with pLDDT > 80. | S36 |
| **Fig. S28. Conformational clustering and sequence features for the zero-insertion design.** | S37 |
| **Fig. S29. Conformational clustering and sequence features for 1-deletion.** | S38 |
| **Fig. S30.** **Conformational clustering and sequence features for 2-deletions.** | S39 |
| **Fig. S31. Conformational clustering and sequence features for 1-insertion.** | S40 |
| **Fig. S32. Conformational clustering and sequence features for 3-insertions.** | S41 |
| **Fig. S33. Conformational clustering and sequence features for 4-insertions.** | S42 |
| **Fig. S34. Conformational clustering and sequence features for 2-insertions, 5 repeats.** | S43 |
| **Fig. S35. Conformational clustering and sequence features for 2-insertions, 6 repeats.** | S44 |
| **Table S1. Top 10 cluster centroid structures across all repeat sizes.** | S45 |
| **Table S2.** Data collection and refinement statistics of Heliscrew4. | S46 |

### **Generation and Structure Prediction of Random Repeat Sequences**

To systematically investigate how repeat unit length influences structural diversity and designability, we generated random amino acid sequences with repeat lengths ranging from 5 to 120 residues. Each random repeat was constructed from an independently sampled amino acid sequence of length n (the repeat unit length), which was then **repeated iteratively to achieve a total chain length of approximately 120 amino acids.**

For example, a 5-residue repeat unit was concatenated 24 times (5 × 24 = 120), whereas a 10-residue repeat unit was repeated 12 times (10 × 12 = 120). When the repeat length did not evenly divide 120, the repeat was truncated or extended slightly to maintain a consistent total length. The repeat lengths analyzed included: **5, 6, 7, 8, 9, 10, 11, 12, 13, 15, 17, 20, 24, 30, 40, 60, and 120.**

| **Repeat Length (n)** | **Number of Repeats** | **Total Length (aa)** |
| --- | --- | --- |
| 5 | 24 | 120 |
| 6 | 20 | 120 |
| 7 | 17 | 119 |
| 8 | 15 | 120 |
| 9 | 13 | 117 |
| 10 | 12 | 120 |
| 11 | 11 | 121 |
| 12 | 10 | 120 |
| 13 | 9 | 117 |
| 15 | 8 | 120 |
| 17 | 7 | 119 |
| 20 | 6 | 120 |
| 24 | 5 | 120 |
| 30 | 4 | 120 |
| 40 | 3 | 120 |
| 60 | 2 | 120 |
| 120 | 1 | 120 |

All amino acid positions within each repeat unit were sampled uniformly from the full 20-amino acid alphabet unless otherwise specified. For repeat lengths up to n = 5, the **entire combinatorial sequence space** was exhaustively sampled, i.e. all $\frac{{20}^{5}}{5}=640,000$possible 5-mer repeat sequences were generated.

However, as the repeat length increased, exhaustive enumeration became computationally infeasible due to the exponential growth of sequence space. Therefore, for repeat lengths greater than 5, we uniformly sampled **approximately 1,000,000 unique sequences** per repeat length.

This approach provided balanced coverage across short and long repeat motifs while maintaining computational tractability. The sampling ensured that each repeat length contributed an equivalent number of structures to the downstream structural prediction and clustering pipeline, thereby minimizing statistical bias toward shorter repeats.

All sequences were submitted to **RaptorX**, a deep-learning–based protein structure prediction server, for tertiary structure modeling. Each structure was evaluated by its **predicted local distance difference test (pLDDT)** confidence score, a per-residue reliability metric ranging from 0 to 100.

Predicted models with **mean pLDDT ≥ 90** were considered high-confidence and retained for further analysis. For 120mers, models with mean pLDDT **≥ 80** were retained. Models below this threshold were discarded. The remaining set of high-confidence structures constituted the **final dataset for clustering and cluster analysis** (described above), ensuring that subsequent analyses were based on well-folded, confidently predicted structural ensembles.

**Primitive amino acid structure generation**

To investigate potential evolutionary constraints on repeat protein formation, we repeated a subset of our structure generation and clustering analyses using a restricted alphabet of **10 primitive amino acids**, hereafter referred to as the **“early 10” set: G, A, S, T, D, E, I, L, P, and V.**

This subset was selected based on prior hypotheses regarding prebiotic amino acid availability and early biosynthetic accessibility, representing a plausible early-stage chemical toolkit for protein evolution. In addition to providing evolutionary insight, limiting the amino acid alphabet significantly reduced the combinatorial complexity of the sequence space, thereby facilitating deeper sampling coverage at a fixed computational cost.

Random repeat sequences were generated using the same stochastic framework employed for the full 20-amino acid set. However, residue selection at each position was restricted to the “early 10” alphabet. For an *n*-residue repeat unit, the total possible sequence space is given by

$$N=k^{n}$$

where *k* represents the alphabet size. Reducing *k* from 20 to 10 thus decreases the combinatorial search space by a factor of

$${(\frac{10}{20})}^{n}=2^{-n}$$

This exponential reduction allows proportionally greater sampling coverage per computational batch. For example, for a 10-residue repeat unit, restricting to 10 amino acids effectively doubles the proportion of sequence space explored for a fixed number of sampled sequences. In practical terms, even modest sampling (e.g., 250,000 generated sequences per repeat length) results in much higher fractional coverage of the total possible sequence space compared to the standard 20-amino acid library. For instance, performing 250,000 samples for each repeat length between 5–10 covers several orders of magnitude more combinations relative to 20-amino acid alphabet sampling.

### **Cluster Analysis Pipeline**

We performed a systematic structural clustering and sequence analysis pipeline to investigate the folding behavior of randomly generated repeat protein sequences. Our goal was to determine how these sequences converge toward stable structural motifs and quantify the degree of designability within each cluster. The strategy consisted of (1) clustering similar structures based on geometric alignment, (2) extract consensus sequences from aligned cluster centroids, and (3) analyzing their conformational preferences and conserved sequence features.

For each repeat length population of predicted repeat structures, the pipeline began by randomly selecting a single model to serve as the cluster centroid. Structural comparisons between this seed and all other predicted structures were performed using TM-align (version 2020), employing both the TM-score and RMSD metrics to evaluate structural similarity. Structures with a root-mean-square deviation (RMSD) ≤ 1.2 Å and a TM-score ≥ 0.7 relative to the cluster centroid were considered members of the same cluster. Structures meeting these criteria were assigned to that cluster, and a new random unclustered structure was selected as the next cluster centroid. This process was repeated iteratively until the entire population of structures had been clustered.

Following clustering, we computed the total number of clusters, the number of members and unique sequences in each cluster to calculate designability. In addition, we determined the cumulative number of clusters required to capture 99% of all predicted structures in a given repeat population, which was used to quantify overall fold diversity.

We then carried out sequence alignment for each cluster by parsing the TM-align output logs and reconstructing the aligned amino acid registers. For each aligned set, we computed a consensus sequence by taking the most frequently observed residue at each aligned position. This was achieved via a custom Python code that performed a column-wise majority rule calculation across the aligned sequences. Each consensus sequence was written to a FASTA file in the corresponding cluster directory and served as a representative model for downstream analyses. These consensus sequences allowed us to compare sequence convergence trends across clusters and relate sequence conservation patterns to fold stability and geometry.

We next performed a detailed backbone torsion angle analysis for each cluster centroid to visualize local conformational preferences. The centroid model was parsed using the Bio.PDB module of Biopython (version 1.83), and the φ and ψ dihedral angles were computed for all residues. We focused on the repeat segment corresponding to the primary structural motif of interest. The resulting φ/ψ values were projected onto an APBLE background map, which categorizes conformational space into α-helix (A), β-sheet (B), left-handed helix (L), and extended (E) regions. The visualization enabled us to directly assess whether repeat units adopted α-helical, β-sheet, or irregular backbone conformations.

To visualize residue conservation within each aligned sequence ensemble, we constructed sequence logos using Logomaker (version 0.8). The sequence logos were generated from position frequency matrices derived from the aligned sequences and converted to information content (bits) per position. We adopted a consistent color scheme reflecting residue physicochemical categories to improve interpretability.

For helical repeat motifs, we performed an analogous analysis, but with additional annotation to capture periodic features of the helix. In these cases, residues were indexed according to heptad notation (A–G), reflecting the seven-residue periodicity typical of coiled-coil helices. We also computed solvent-accessible surface area (SASA) values for each position using structural models of the cluster centroids, which were then displayed above each register position in the sequence logo. This visualization provided a direct link between conservation patterns and solvent exposure, highlighting which heptad positions were buried versus exposed in the helical bundle. The plotting routine used for these helical logos is illustrated below:

In both cases, the resulting images provided quantitative visualization of residue conservation patterns and positional variability, allowing us to compare the sequence features of highly designable clusters against less populated structural motifs. The incorporation of SASA information further revealed that conserved residues were often located at buried positions in the helical core, whereas solvent-exposed registers exhibited greater variability across aligned sequences.

The amino acid coloring scheme employed throughout this analysis is summarized in the following table:

| **Amino Acid** | **Category** | **Color (Hex)** |
| --- | --- | --- |
| R, K, D, E | Charged | #FF5500 |
| Q, N, H | Polar uncharged | #FF8844 |
| S, T, C | Small polar | #FFBB88 |
| A | Hydrophobic | #88BBFF |
| Y, W, F | Aromatic | #4488FF |
| I, V, L, M | Hydrophobic | #0044FF |
| G | Special (Glycine) | #000000 |
| P | Special (Proline) | #FFD700 |

All computational analyses were performed using Python 3.10 in a Conda environment on macOS 14.6.1. Key packages included numpy (v1.26), pandas (v2.2), matplotlib (v3.9), biopython (v1.83), and logomaker (v0.8). Structural alignment was performed using TM-align (v2020). All scripts were executed serially on an 8-core CPU workstation with fixed random seeds to ensure reproducibility. Intermediate files, including aligned FASTA sequences, consensus outputs, torsion angle tables, and sequence logo images, were retained for full traceability of the computational workflow.

**Structural Database Searches**

In order to place our *de novo* cluster centroid structures in the context of known protein space, we performed a structural homology search against a nonredundant PDB set and applied domain-level annotation via CATH coverage. For each cluster centroid, we aligned the structure to entries in the **PDB60** dataset (a subset filtered at ≤ 60 % sequence homology) using **TM-align** (v2020). We collected all PDB targets yielding **TM-score ≥ 0.5** when aligned to the centroid, then ranked these hits by descending TM-score. The top hits (usually > 5) were retained for further inspection and annotation.

To complement this, we applied **CATHcover** (a publicly available method for mapping structures to CATH domain classifications) to identify possible domain-level matches even when global structural similarity was weak. CATHcover takes as input a query structure and finds the minimal set of CATH domains whose combination best “covers” the query’s topology. By combining the TM-align ranking and CATHcover domain mapping, we assigned likely homologous domains (or architectural analogs) to many of our centroid structures, thereby relating our cluster folds to known structural families in the PDB or CATH hierarchy.

In practice, we prefiltered candidate PDBs by chain length (within ± 20 residues of 120 aa) and by structural class (where available) to reduce computational burden. We then ran TM-align only on that filtered subset. The hits passing the TM-score threshold were annotated with PDB IDs, chain identifiers, TM-score, and RMSD, and incorporated into the supplemental table of centroid–PDB matches.

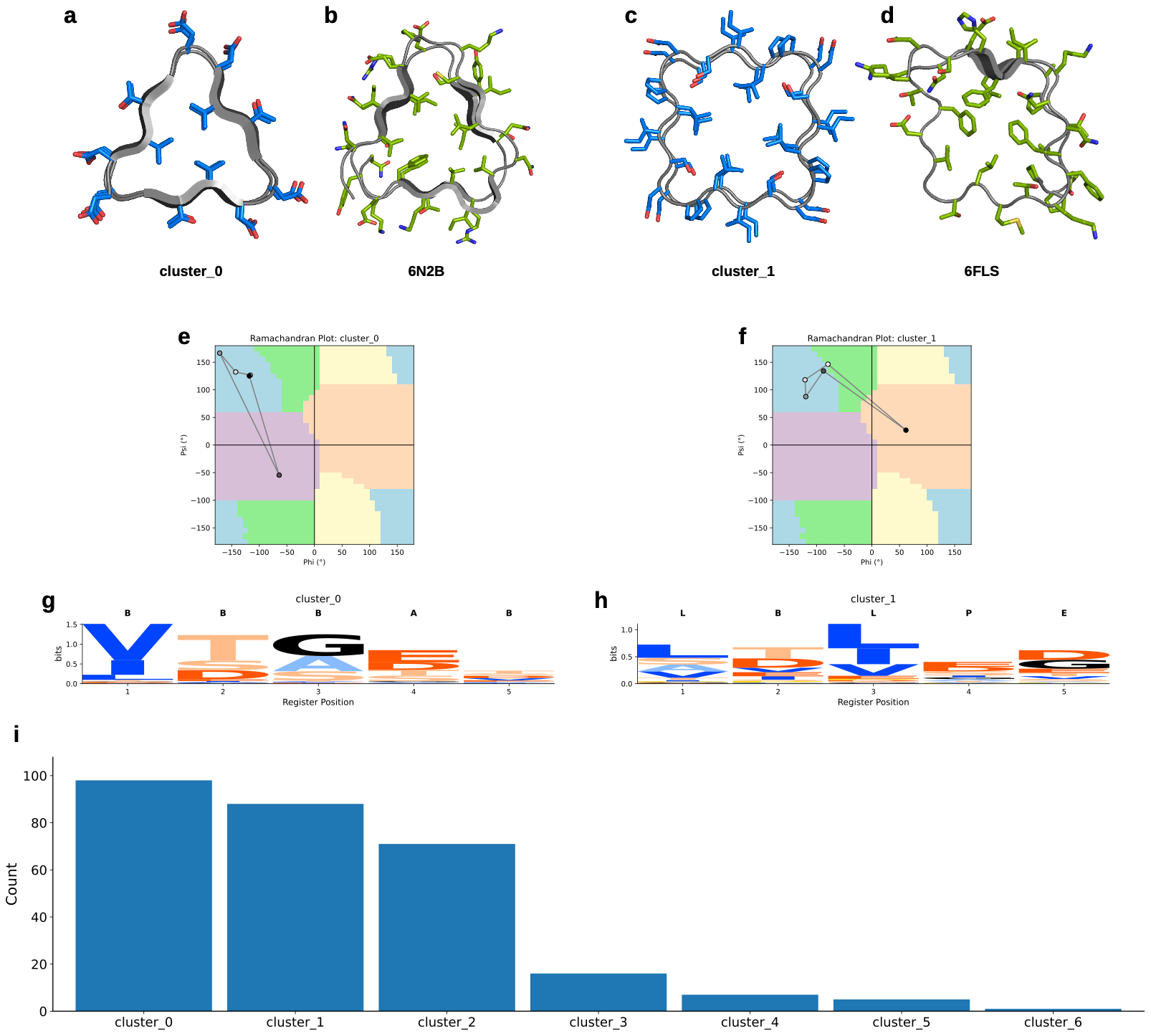

**Fig. S1.** Conformational clustering and sequence features of pentameric repeat proteins generated from “primitive” amino acid set. (A-D) Representative structures from cluster_0 and cluster_1 compared with nearest PDB analogues (6N2B, 6FLS). (E, F) Ramachandran plots showing backbone angle distributions for clusters 0 and 1. (G, H) Sequence logos for each cluster, illustrating register-specific residue preferences. (I) Cluster size distribution for all designs.

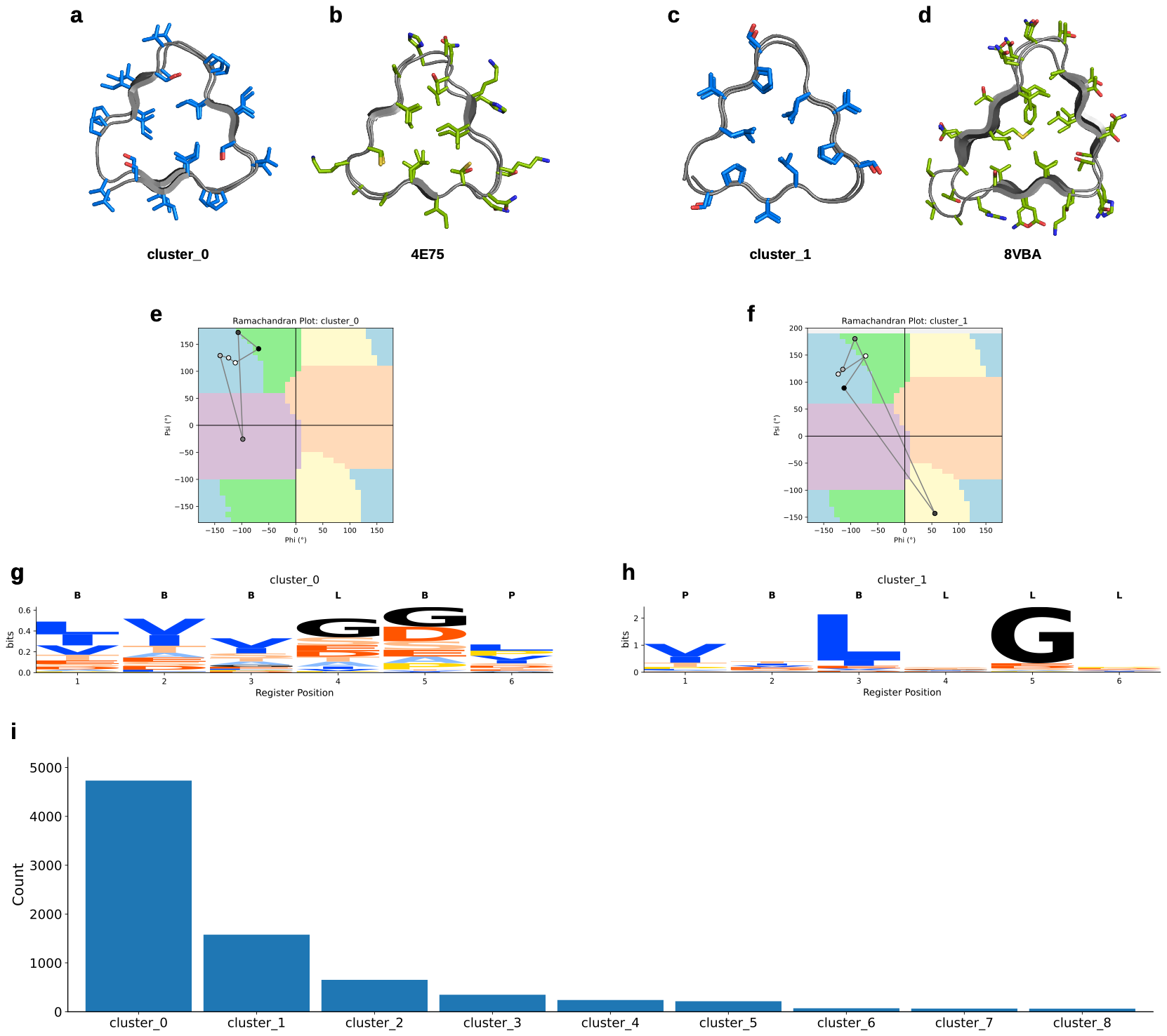

**Fig. S2.** Conformational clustering and sequence features of hexameric repeat proteins generated from “primitive” amino acid set. (A–D) Representative structures from cluster_0 and cluster_1 compared with their nearest PDB analogues (4E75, 8VBA). (E, F) Ramachandran plots showing backbone angle distributions for clusters 0 and 1. (G, H) Sequence logos for each cluster, illustrating register-specific residue preferences across the 6-mer repeat. (I) Cluster size distribution for all designs.

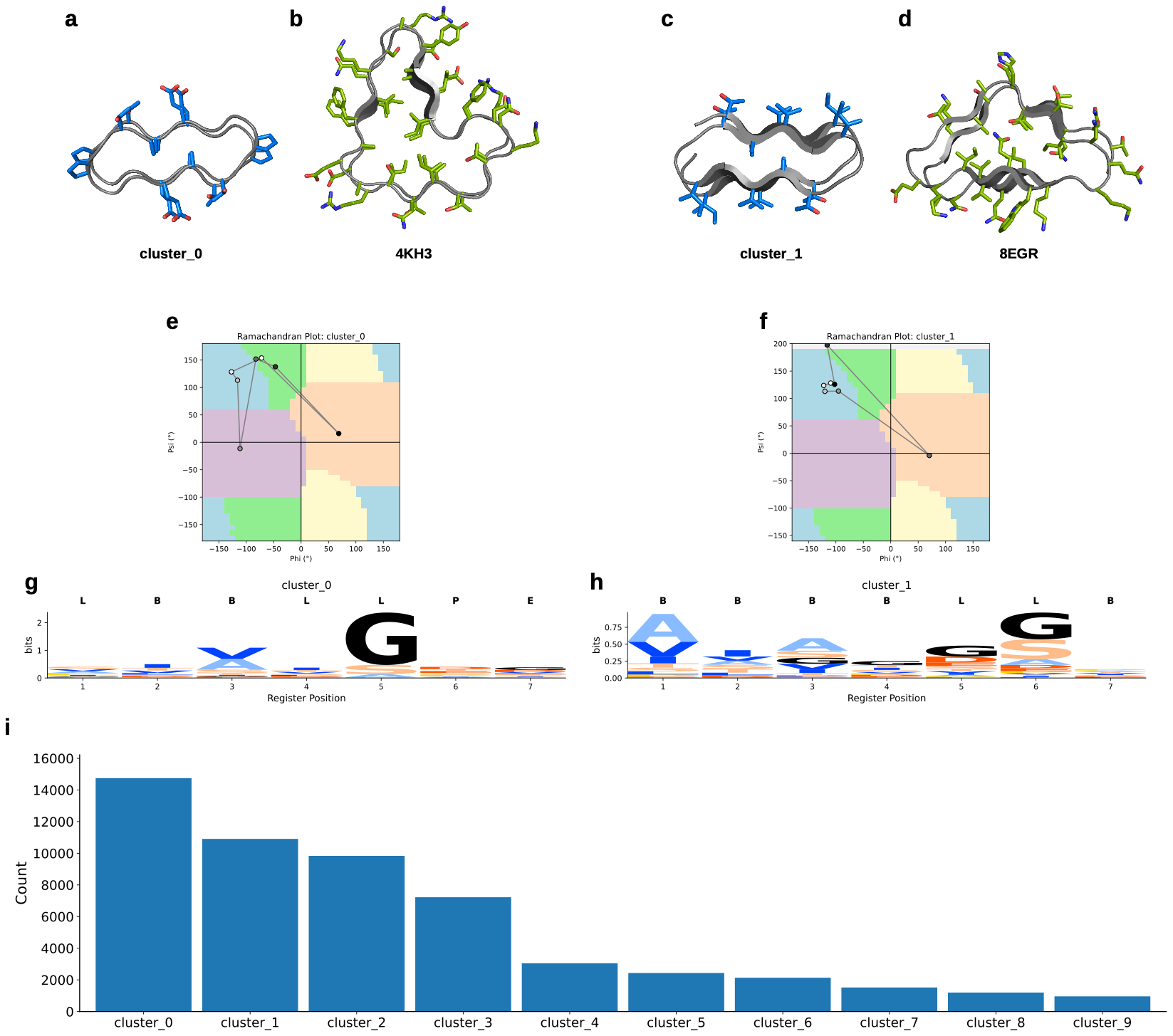

**Fig. S3. Conformational clustering and sequence features of designed 7-mer repeat proteins generated from the “primitive” amino acid set.** (A–D) Representative structures from cluster_0 and cluster_1 compared with their nearest PDB analogues (4KH3, 8EGR). (E, F) Ramachandran plots showing backbone angle distributions for clusters 0 and 1. (G, H) Sequence logos for each cluster, illustrating register-specific residue preferences across the 7-mer repeat. (I) Cluster size distribution for all designs.

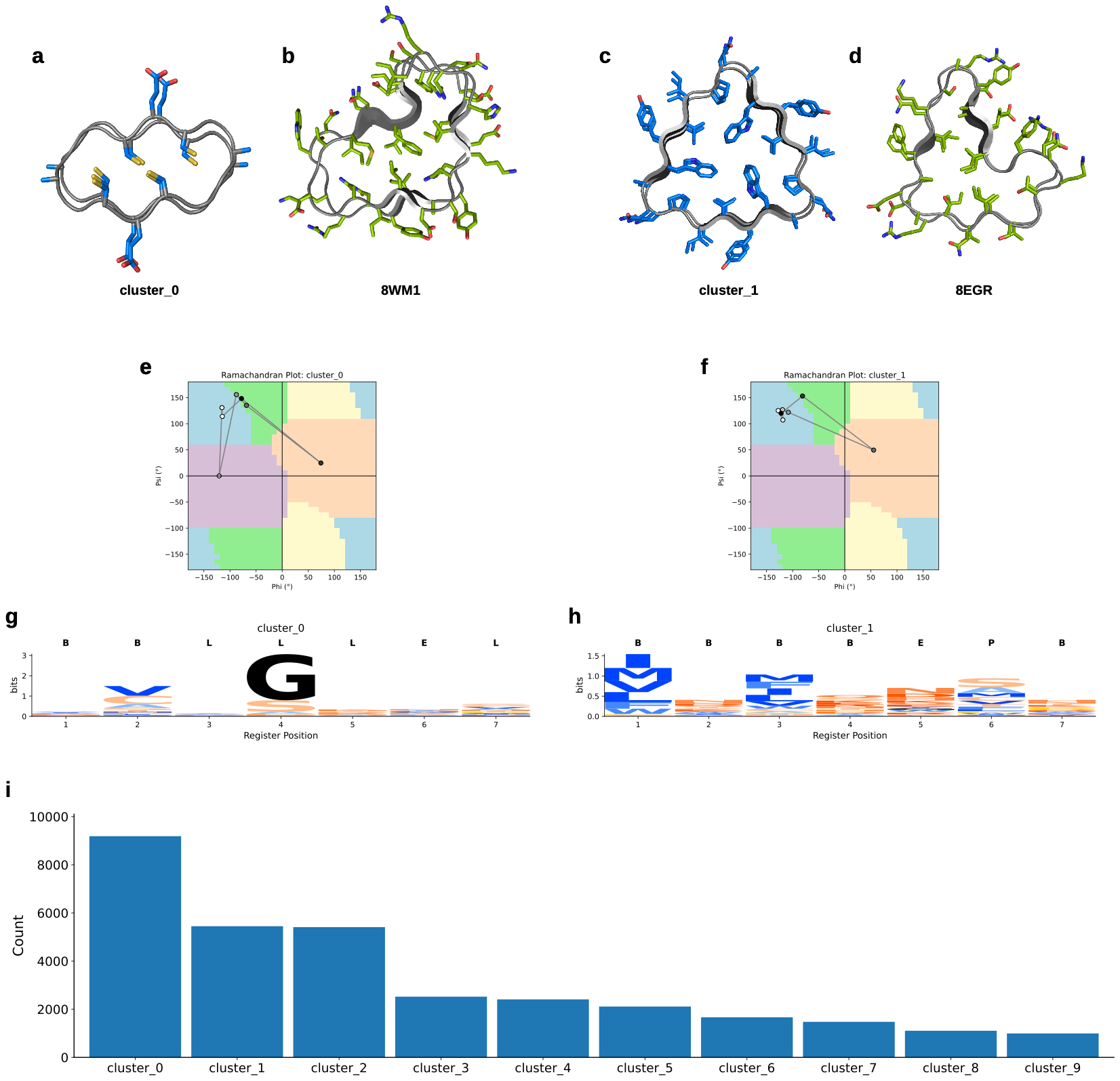

**Fig. S4. Conformational clustering and sequence features of designed 7-mer repeat proteins generated from the full 20 canonical amino acid set.** (A–D) Representative structures from cluster_0 and cluster_1 compared with their nearest PDB analogues (8WM1, 8EGR). (E, F) Ramachandran plots showing backbone angle distributions for clusters 0 and 1. (G, H) Sequence logos for each cluster, illustrating register-specific residue preferences across the 7-mer repeat. (I) Cluster size distribution for all designs.

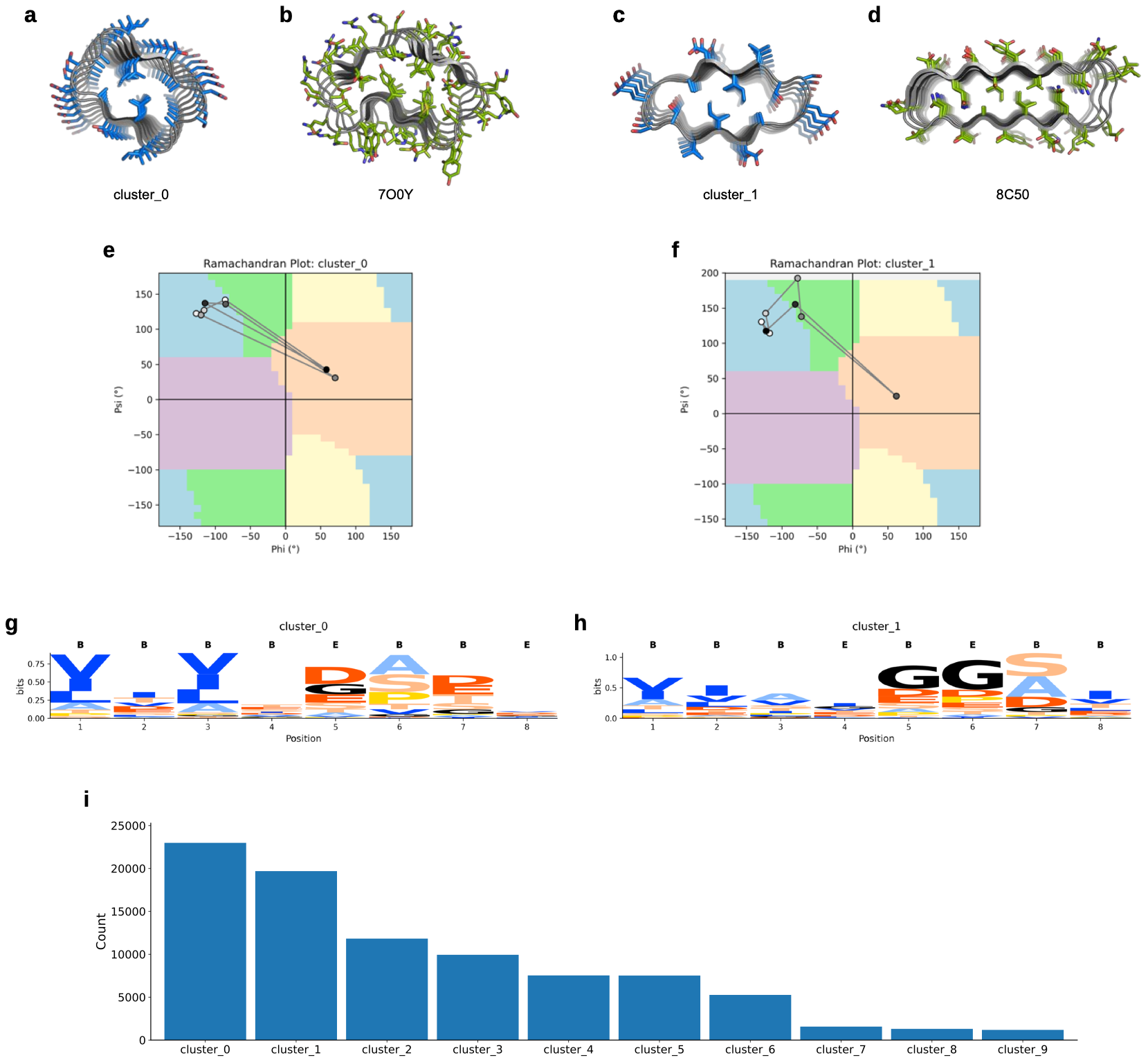

**Fig. S5. Conformational clustering and sequence features of designed 8-mer repeat proteins (early 10 set).** (A–D) Representative structures from cluster_0 and cluster_1 compared with their nearest PDB analogues (7OOY, 8C50). (E, F) Ramachandran plots showing backbone angle distributions for clusters 0 and 1. (G, H) Sequence logos for each cluster, illustrating register-specific residue preferences across the 8-mer repeat. (I) Cluster size distribution for all designs.

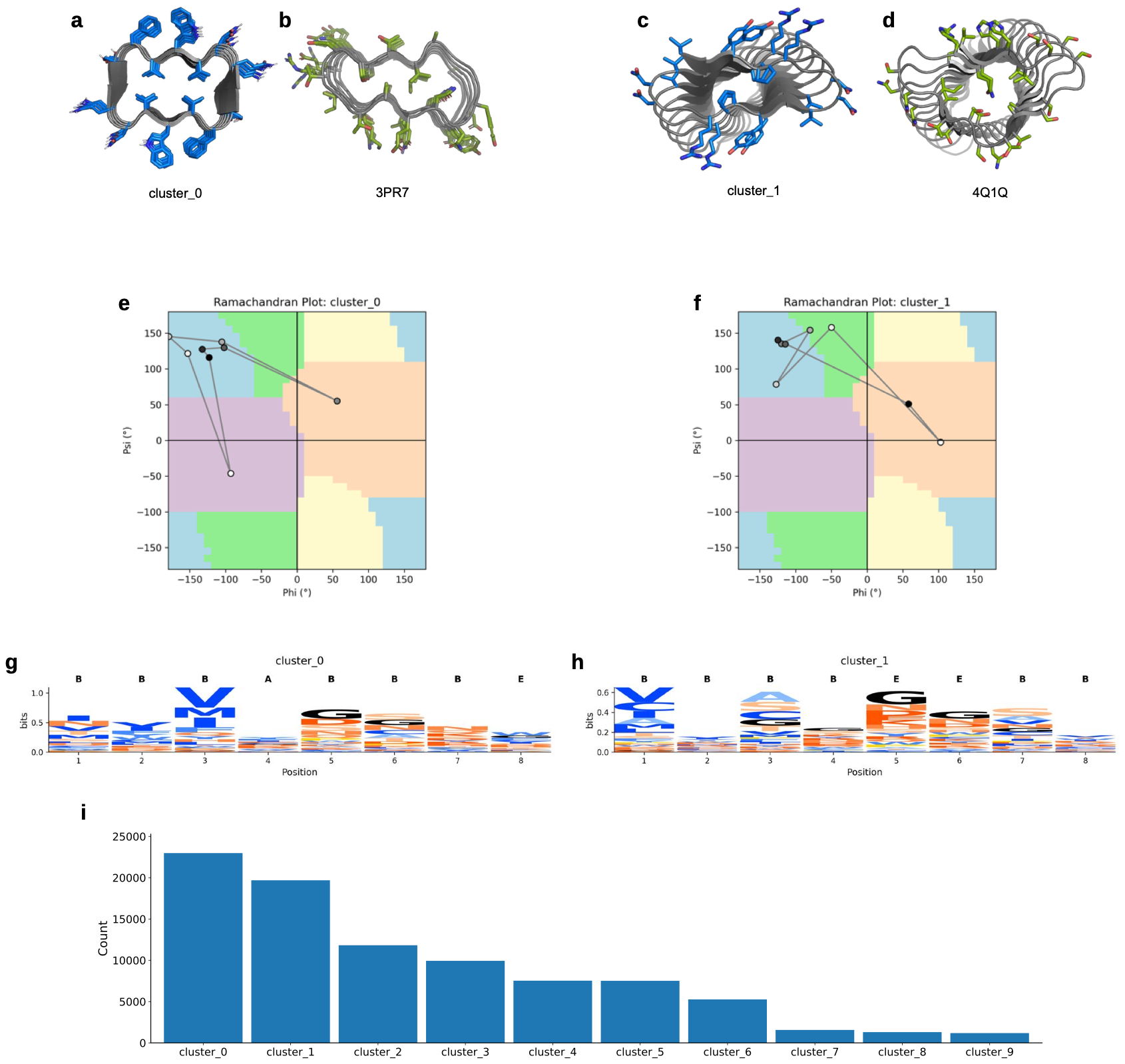

**Fig. S6. Conformational clustering and sequence features of designed 8-mer repeat proteins generated from the full amino acid set.** (A–D) Representative structures from cluster_0 and cluster_1 compared with their nearest PDB analogues (3PR7, 4Q1Q). (E, F) Ramachandran plots showing backbone angle distributions for clusters 0 and 1. (G, H) Sequence logos for each cluster, illustrating register-specific residue preferences across the 8-mer repeat. (I) Cluster size distribution for all designs.

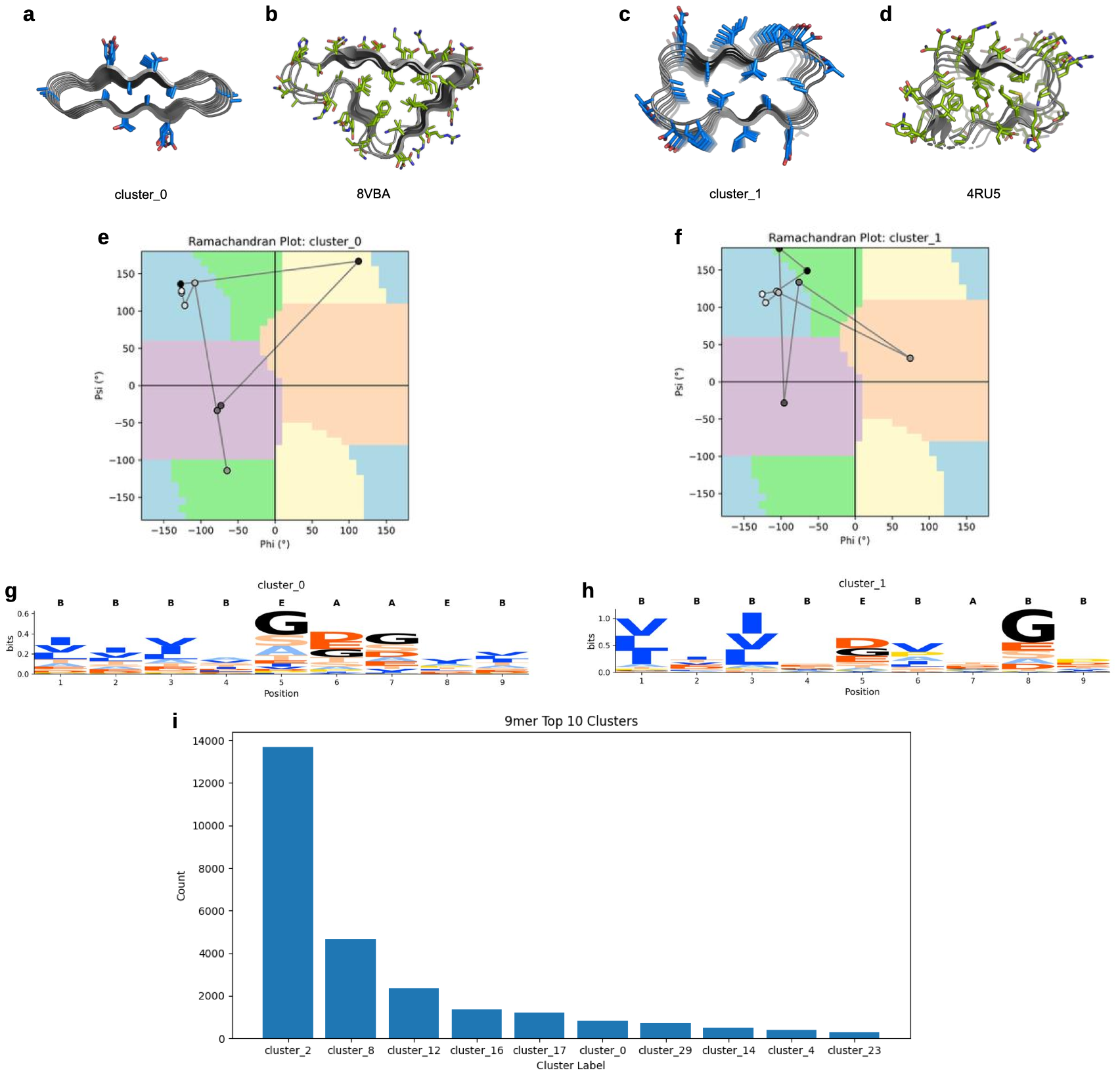

**Fig. S7. Conformational clustering and sequence features of designed 9-mer repeat proteins (early 10 set).** (A–D) Representative structures from cluster_0 and cluster_1 compared with their nearest PDB analogues (8VBA, 4RU5). (E, F) Ramachandran plots showing backbone angle distributions for clusters 0 and 1. (G, H) Sequence logos for each cluster, illustrating register-specific residue preferences across the 9-mer repeat. (I) Cluster size distribution for the top 10 clusters.

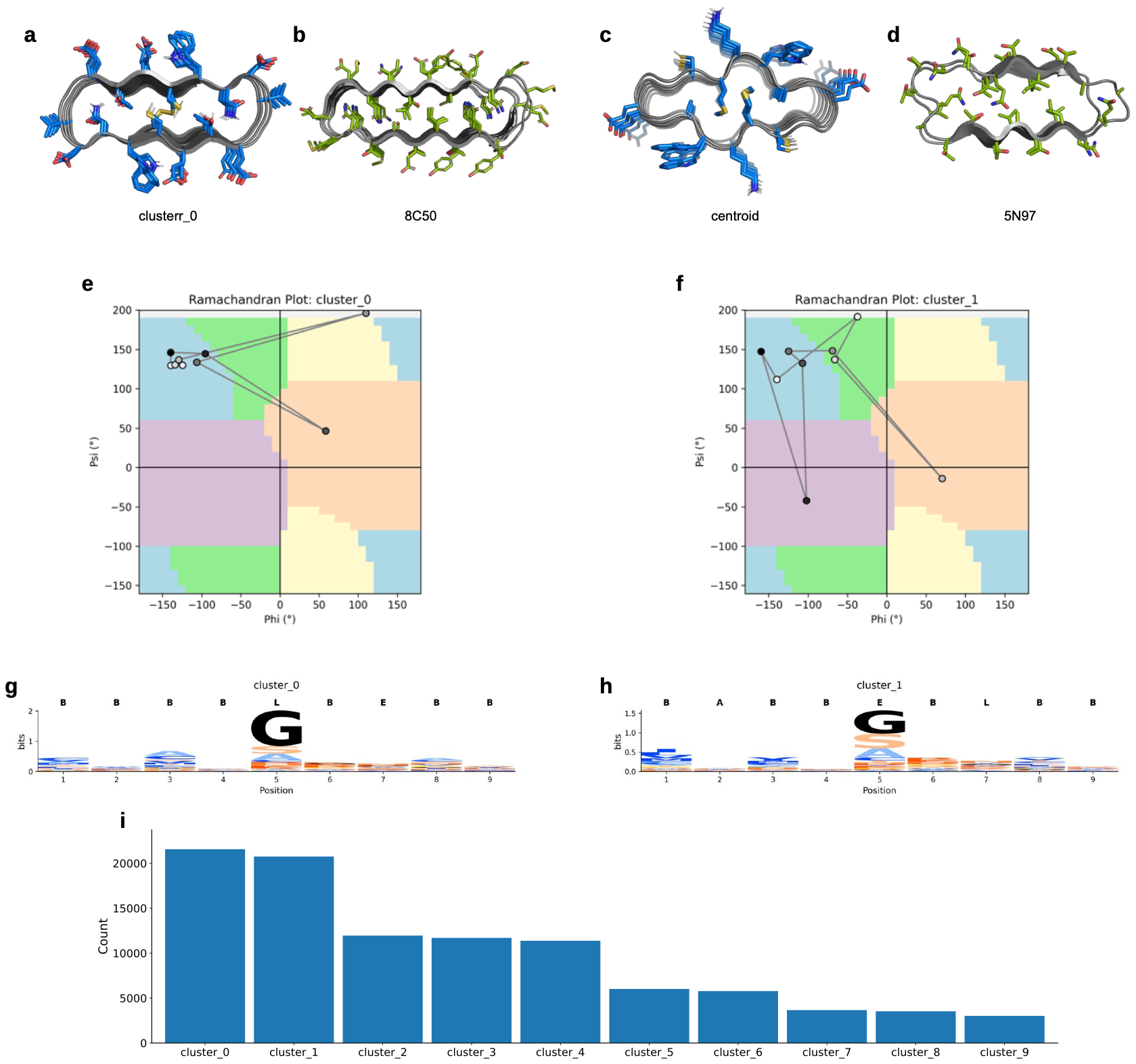

**Fig. S8. Conformational clustering and sequence features of designed 9-mer repeat proteins generated from the full amino acid set.** (A–D) Representative structures from cluster_0 and cluster_1 compared with their nearest PDB analogues (8C50, 5N97). (E, F) Ramachandran plots showing backbone angle distributions for clusters 0 and 1. (G, H) Sequence logos for each cluster, illustrating register-specific residue preferences across the 9-mer repeat. (I) Cluster size distribution for all designs.

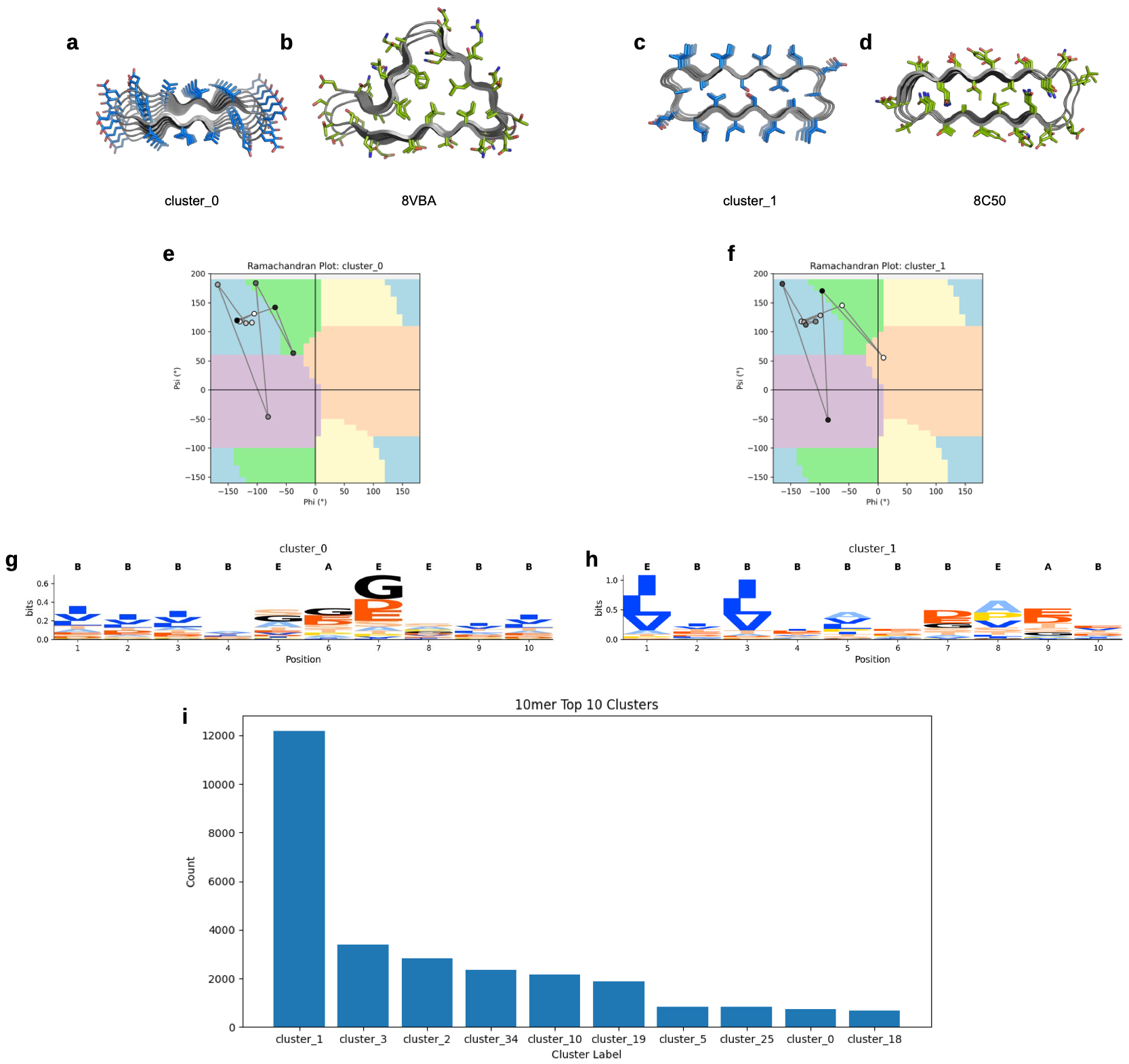

**Fig. S9. Conformational clustering and sequence features of designed 10-mer repeat proteins (early 10 set).** (A–D) Representative structures from cluster_0 and cluster_1 compared with their nearest PDB analogues (8VBA, 8C50). (E, F) Ramachandran plots showing backbone angle distributions for clusters 0 and 1. (G, H) Sequence logos for each cluster, illustrating register-specific residue preferences across the 10-mer repeat. (I) Cluster size distribution for the top 10 clusters.

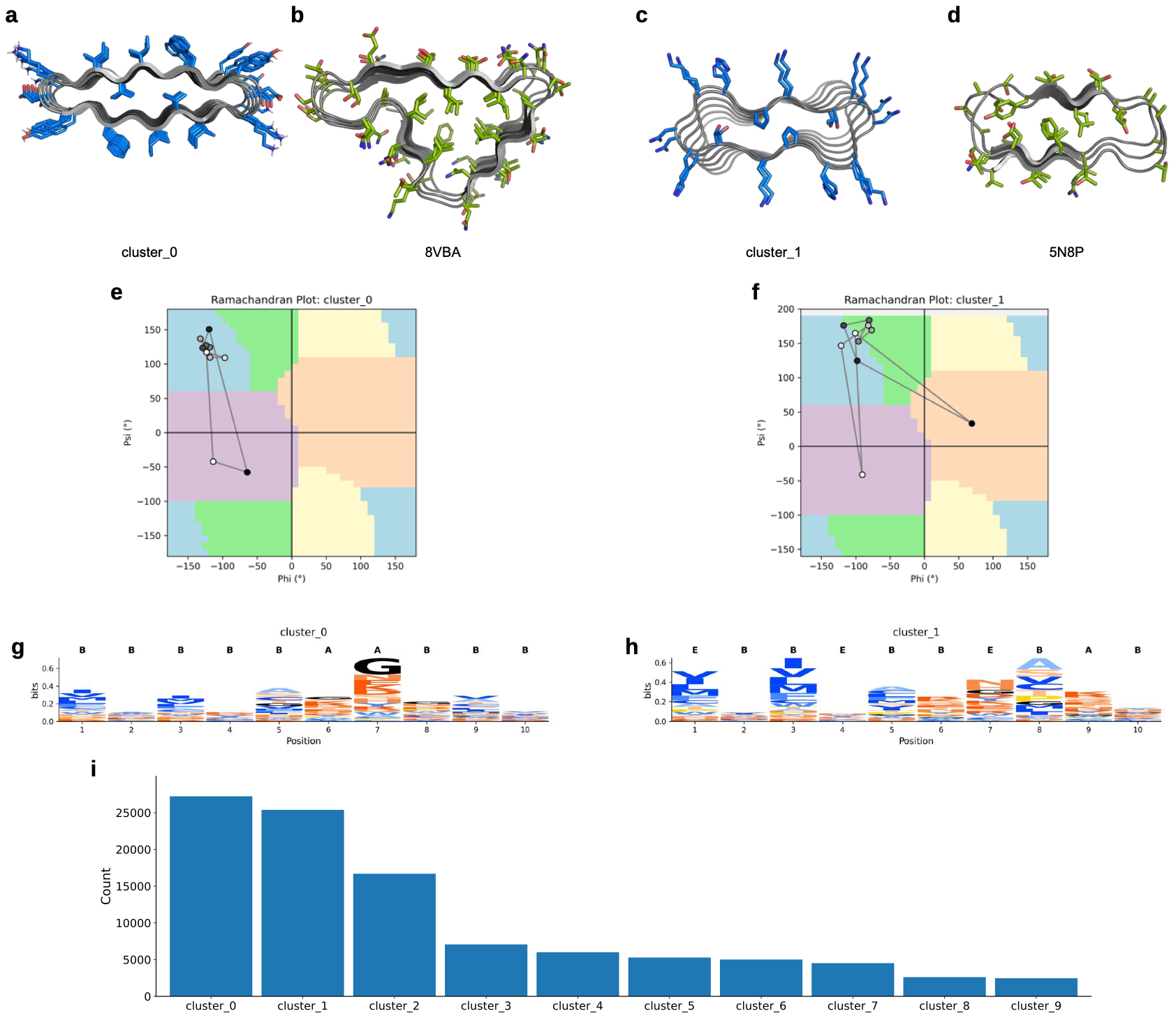

**Fig. S10. Conformational clustering and sequence features of designed 10-mer repeat proteins generated from the full amino acid set. (**A–D) Representative structures from cluster_0 and cluster_1 compared with their nearest PDB analogues (8VBA, 5N8P). (E, F) Ramachandran plots showing backbone angle distributions for clusters 0 and 1. (G, H) Sequence logos for each cluster, illustrating register-specific residue preferences across the 10-mer repeat. (I) Cluster size distribution for all designs.

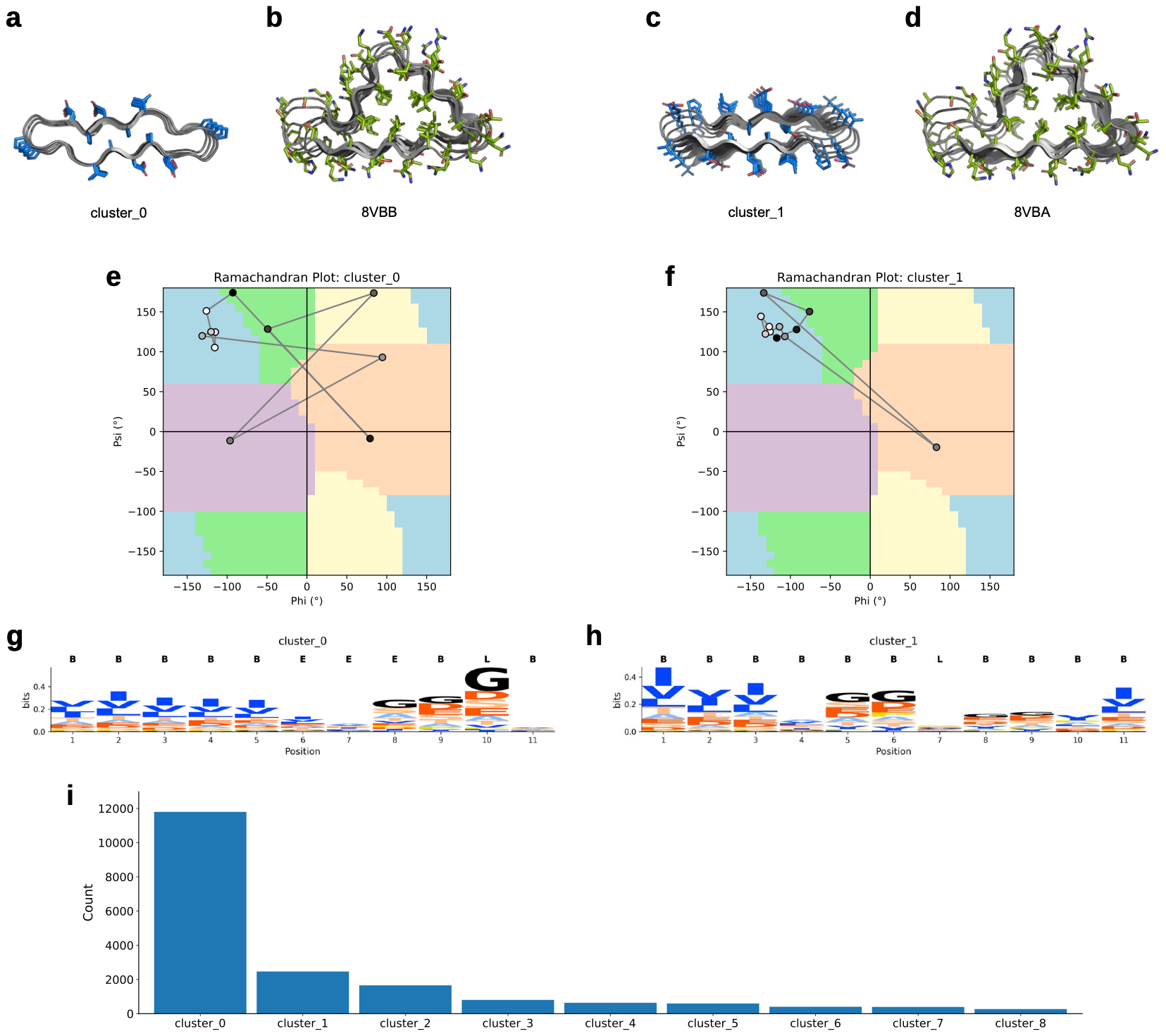

**Fig. S11. Conformational clustering and sequence features of designed 11-mer repeat proteins (early 10 set). (**A–D) Representative structures from cluster_0 and cluster_1 compared with their nearest PDB analogues (8VBB, 8VBA). (E, F) Ramachandran plots showing backbone angle distributions for clusters 0 and 1. (G, H) Sequence logos for each cluster, illustrating register-specific residue preferences across the 11-mer repeat. (I) Cluster size distribution for all designs.

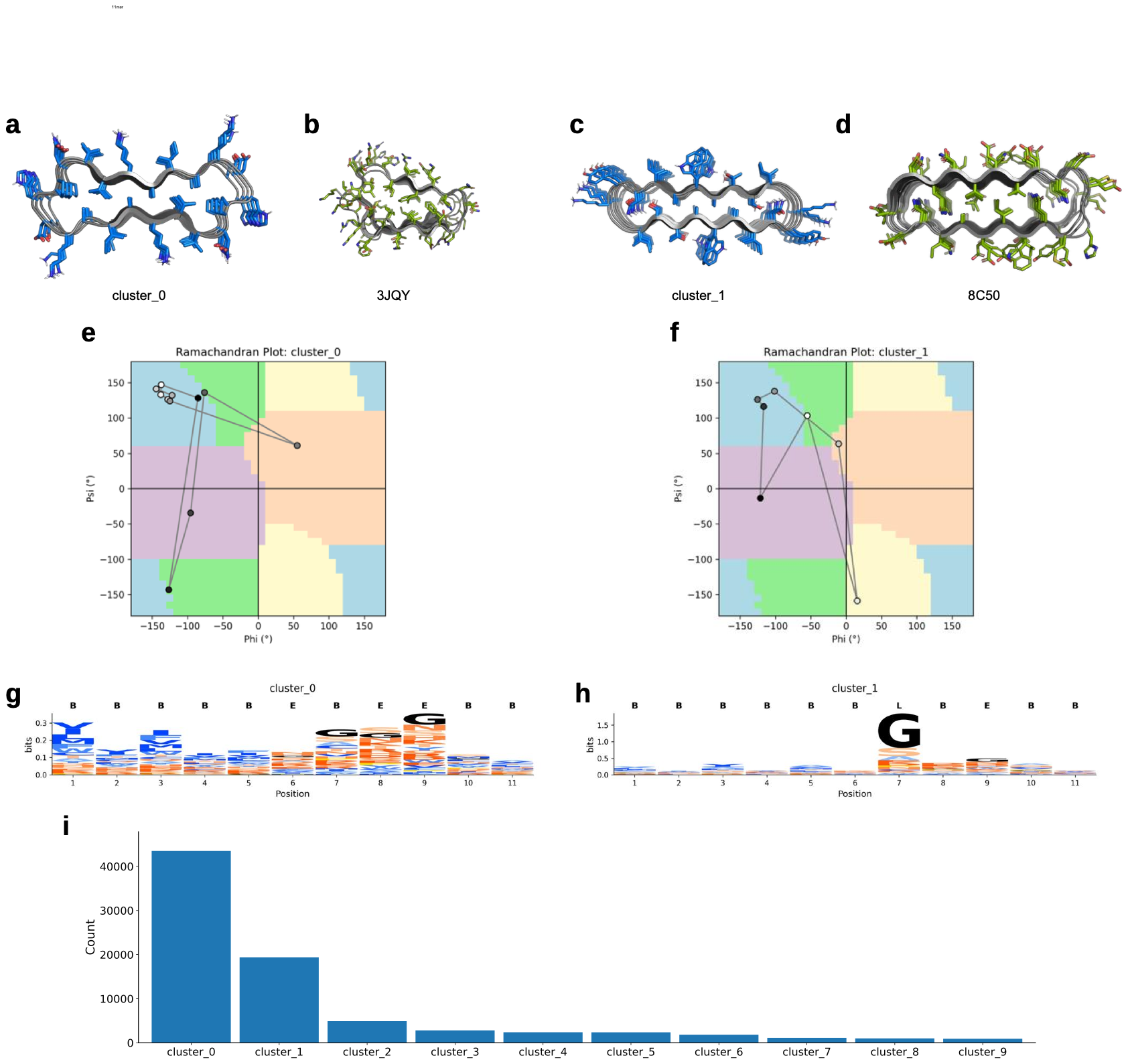

**Fig. S12. Conformational clustering and sequence features of designed 11-mer repeat proteins generated from the full amino acid set. (**A–D) Representative structures from cluster_0 and cluster_1 compared with their nearest PDB analogues (3JQY, 8C50). (E, F) Ramachandran plots showing backbone angle distributions for clusters 0 and 1. (G, H) Sequence logos for each cluster, illustrating register-specific residue preferences across the 11-mer repeat. (I) Cluster size distribution for all designs.

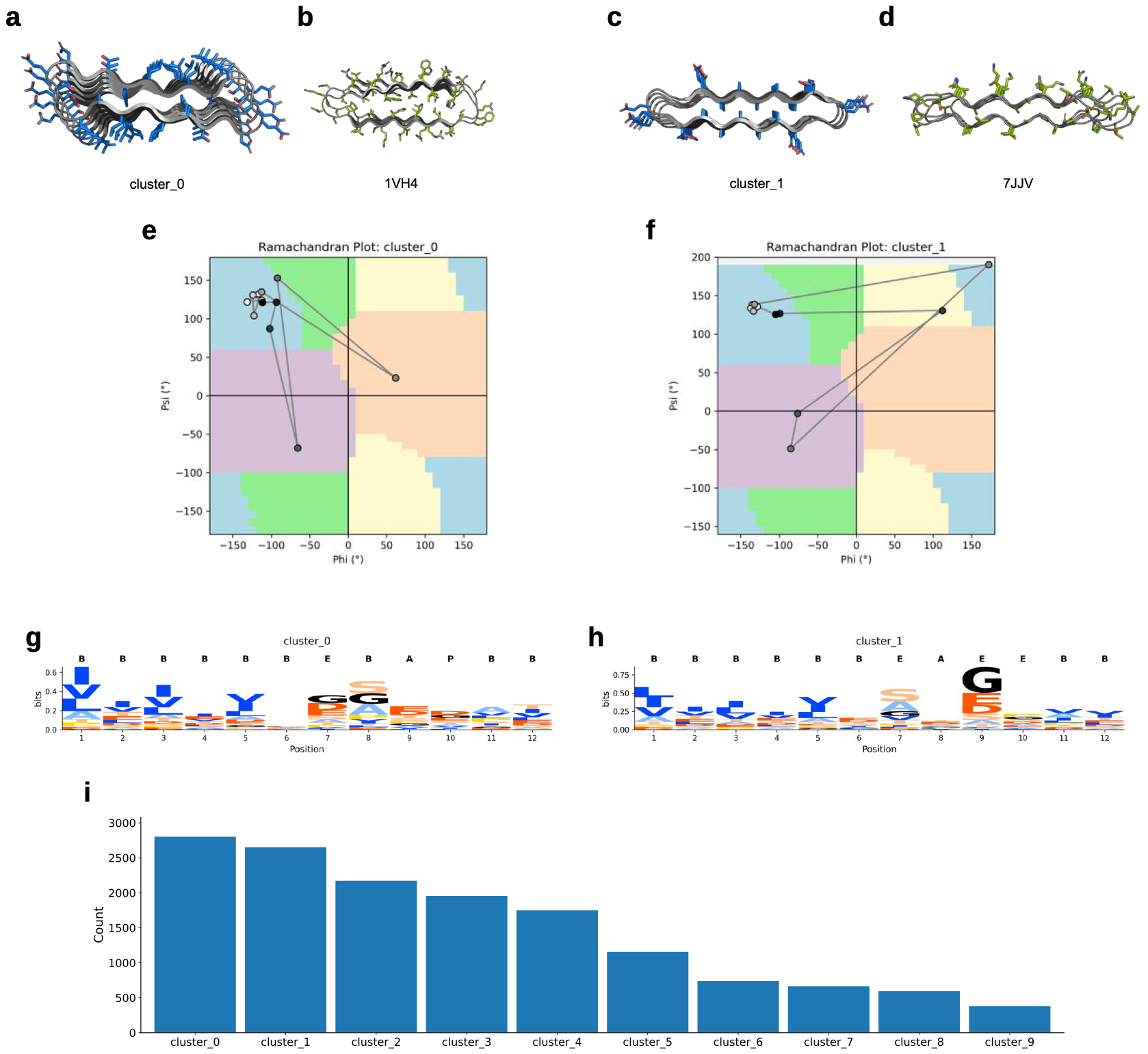

**Fig. S13. Conformational clustering and sequence features of designed 12-mer repeat proteins (early 10 set). (**A–D) Representative structures from cluster_0 and cluster_1 compared with their nearest PDB analogues (1VH4, 7JJV). (E, F) Ramachandran plots showing backbone angle distributions for clusters 0 and 1. (G, H) Sequence logos for each cluster, illustrating register-specific residue preferences across the 12-mer repeat. (I) Cluster size distribution for all designs.

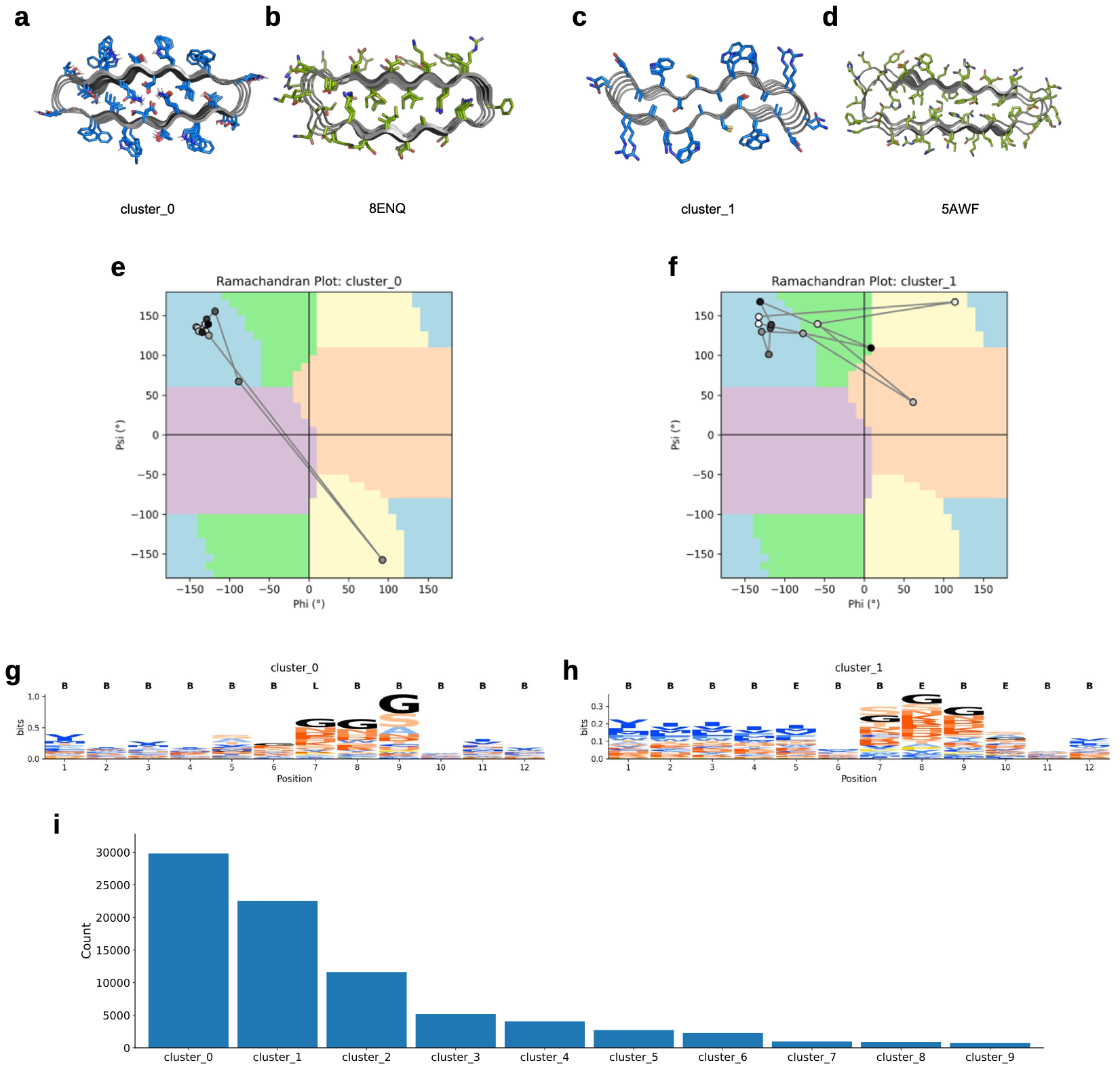

**Fig. S14. Conformational clustering and sequence features of designed 12-mer repeat proteins generated from the full amino acid set. (**A–D) Representative structures from cluster_0 and cluster_1 compared with their nearest PDB analogues (8ENQ, 5AWF). (E, F) Ramachandran plots showing backbone angle distributions for clusters 0 and 1. (G, H) Sequence logos for each cluster, illustrating register-specific residue preferences across the 12-mer repeat. (I) Cluster size distribution for all designs.

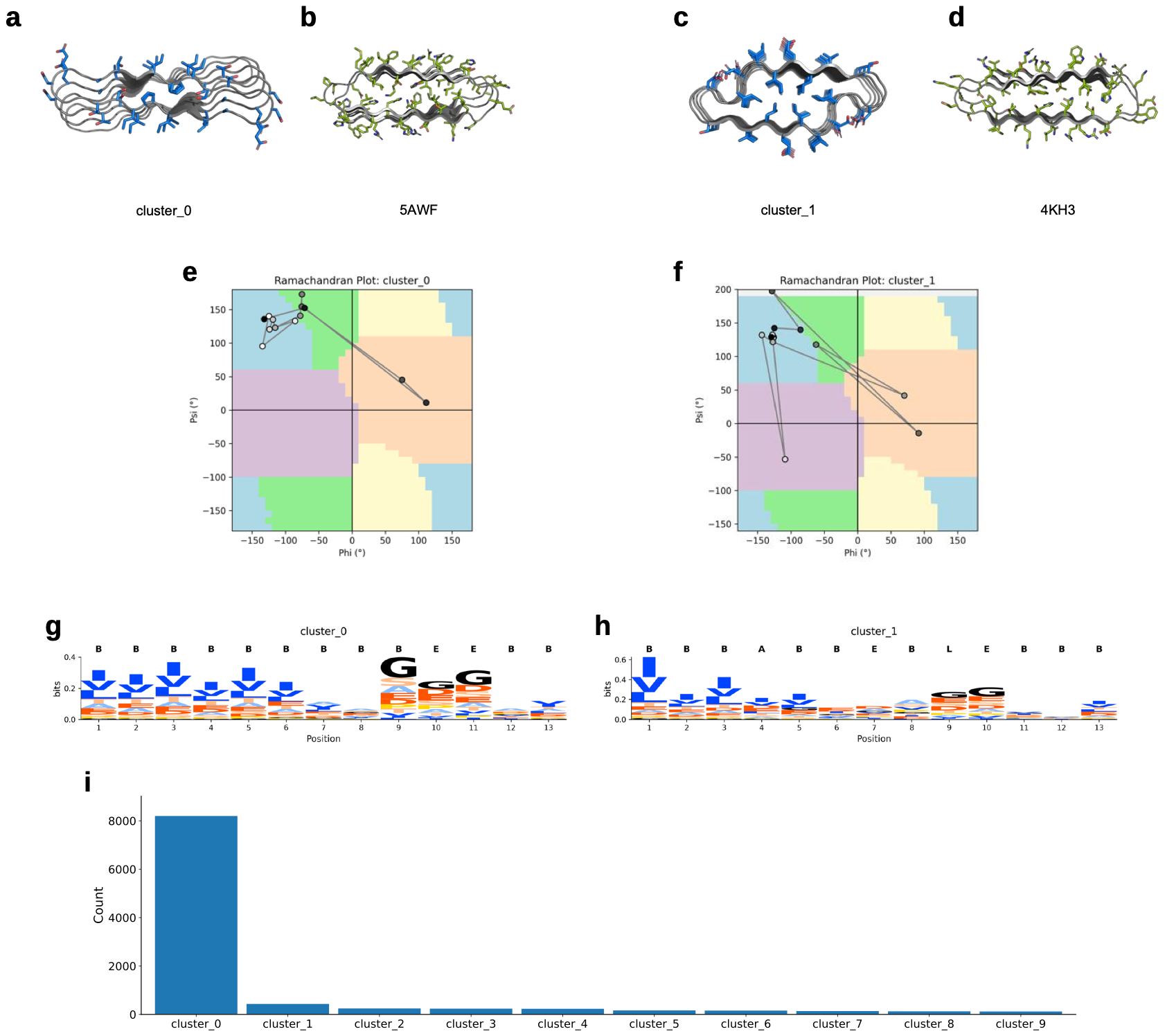

**Fig. S15. Conformational clustering and sequence features of designed 13-mer repeat proteins (early 10 set). (**A–D) Representative structures from cluster_0 and cluster_1 compared with their nearest PDB analogues (5AWF, 4KH3). (E, F) Ramachandran plots showing backbone angle distributions for clusters 0 and 1. (G, H) Sequence logos for each cluster, illustrating register-specific residue preferences across the 13-mer repeat. (I) Cluster size distribution for all designs.

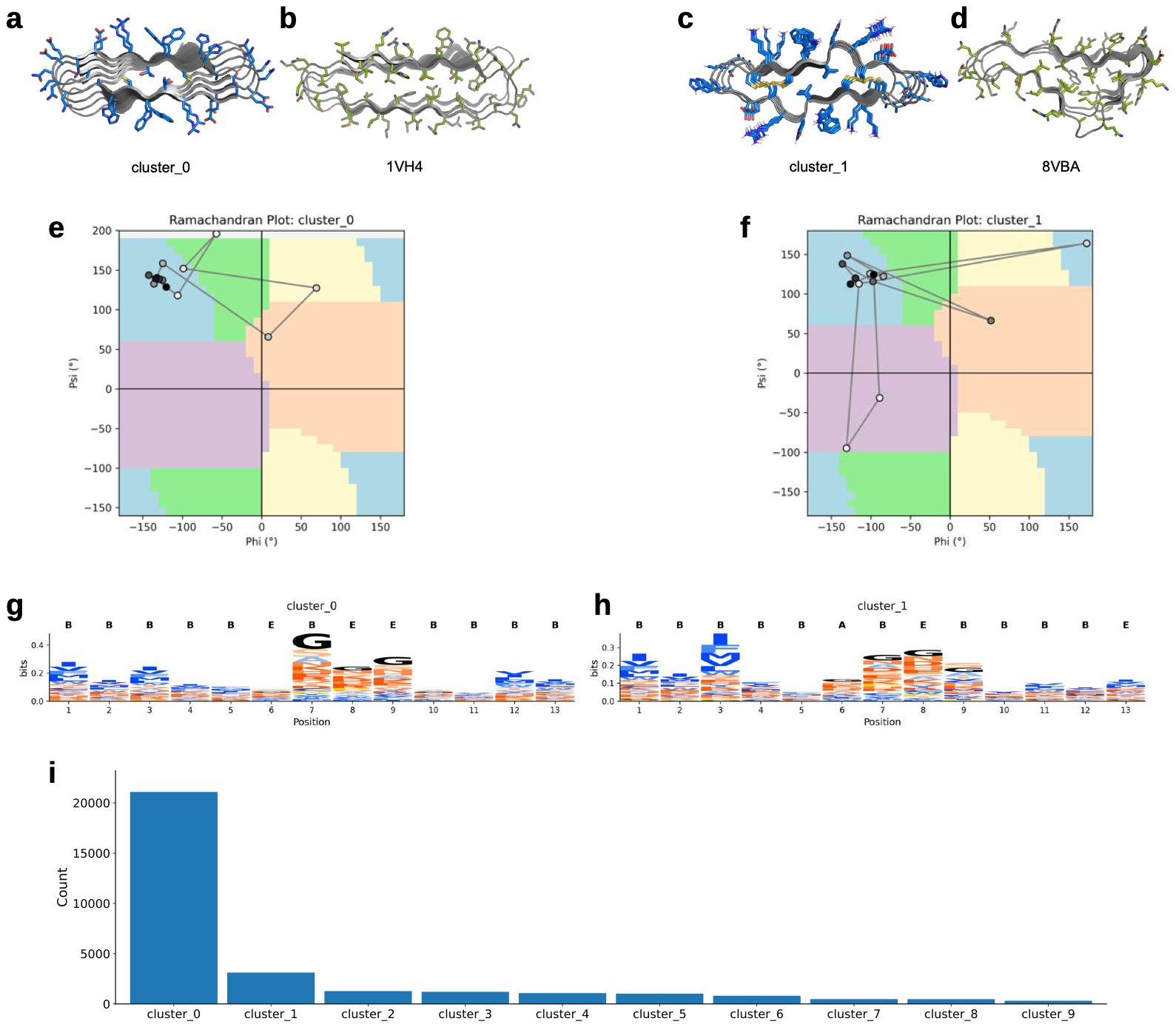

**Fig. S16. Conformational clustering and sequence features of designed 13-mer repeat proteins generated from the full amino acid set. (**A–D) Representative structures from cluster_0 and cluster_1 compared with their nearest PDB analogues (1VH4, 8VBA). (E, F) Ramachandran plots showing backbone angle distributions for clusters 0 and 1. (G, H) Sequence logos for each cluster, illustrating register-specific residue preferences across the 13-mer repeat. (I) Cluster size distribution for all designs.

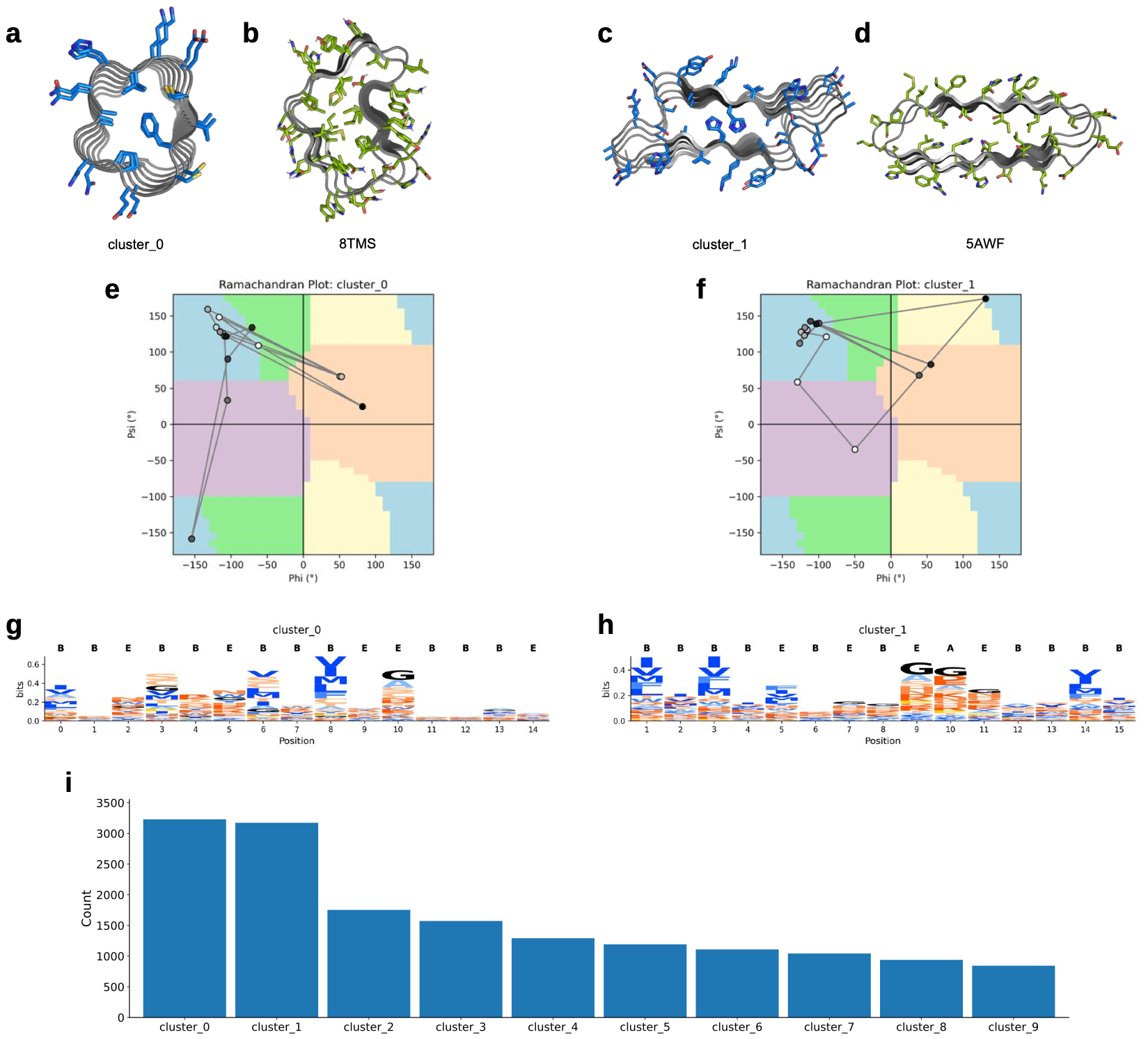

**Fig. S17. Conformational clustering and sequence features of designed 15-mer repeat proteins generated from the full amino acid set. (**A–D) Representative structures from cluster_0 and cluster_1 compared with their nearest PDB analogues (8TMS, 5AWF). (E, F) Ramachandran plots showing backbone angle distributions for clusters 0 and 1. (G, H) Sequence logos for each cluster, illustrating register-specific residue preferences across the 15-mer repeat. (I) Cluster size distribution for all designs.

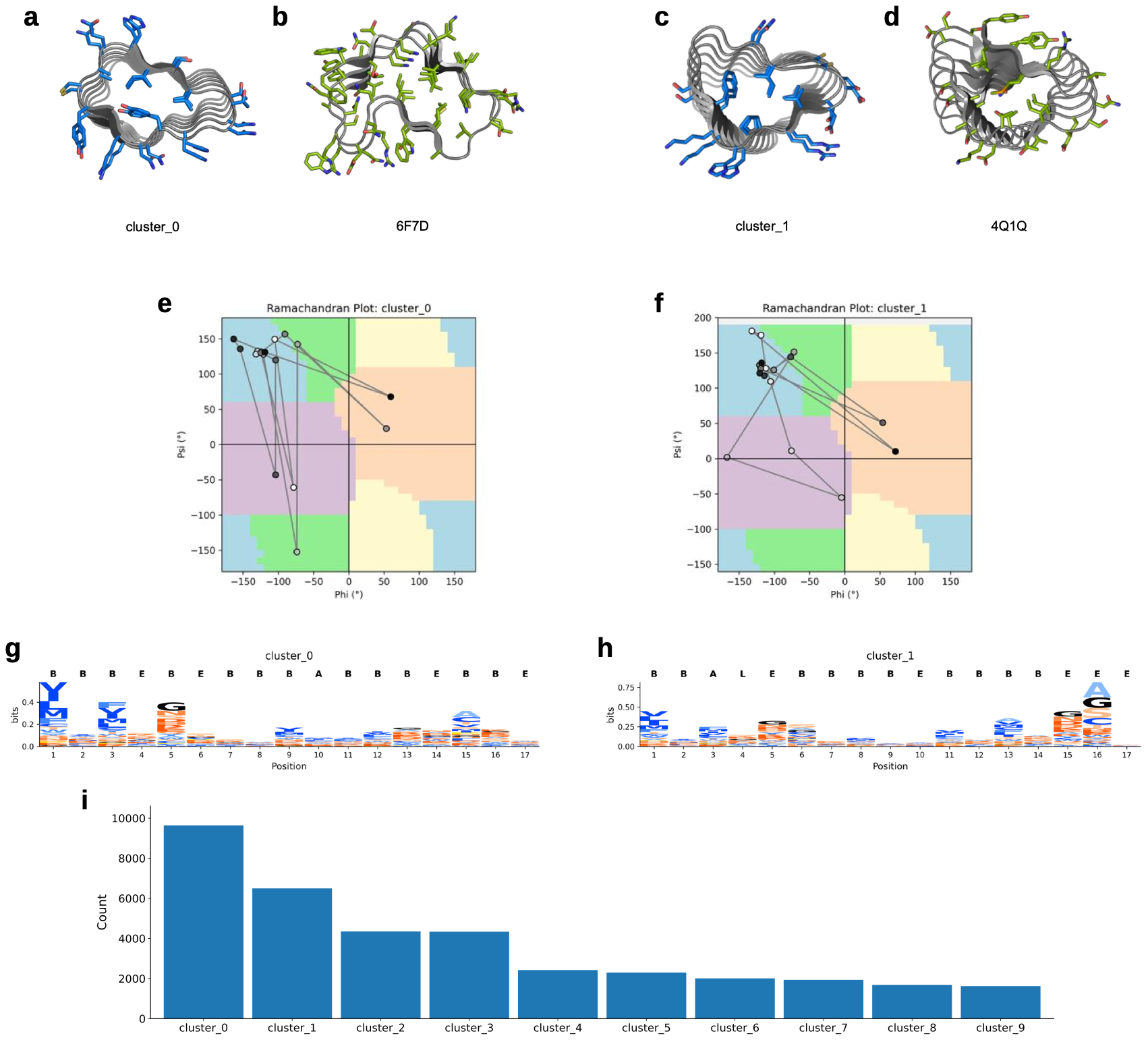

**Fig. S18. Conformational clustering and sequence features of designed 17-mer repeat proteins generated from the full amino acid set. (**A–D) Representative structures from cluster_0 and cluster_1 compared with their nearest PDB analogues (6F7D, 4Q1Q). (E, F) Ramachandran plots showing backbone angle distributions for clusters 0 and 1. (G, H) Sequence logos for each cluster, illustrating register-specific residue preferences across the 17-mer repeat. (I) Cluster size distribution for all designs.

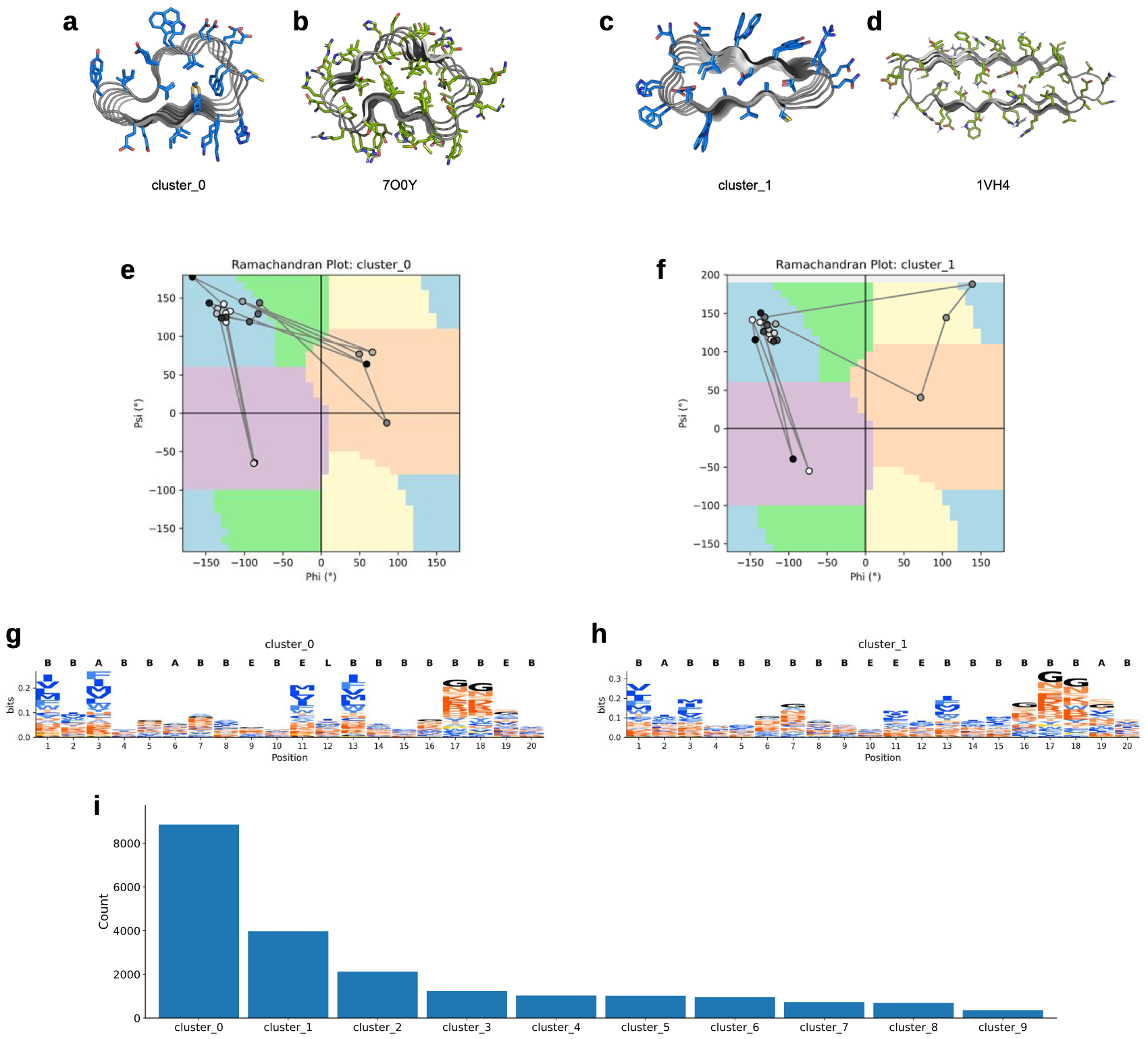

**Fig. S19. Conformational clustering and sequence features of designed 20-mer repeat proteins generated from the full amino acid set. (**A–D) Representative structures from cluster_0 and cluster_1 compared with their nearest PDB analogues (7OQY, 1VH4). (E, F) Ramachandran plots showing backbone angle distributions for clusters 0 and 1. (G, H) Sequence logos for each cluster, illustrating register-specific residue preferences across the 20-mer repeat. (I) Cluster size distribution for all designs.

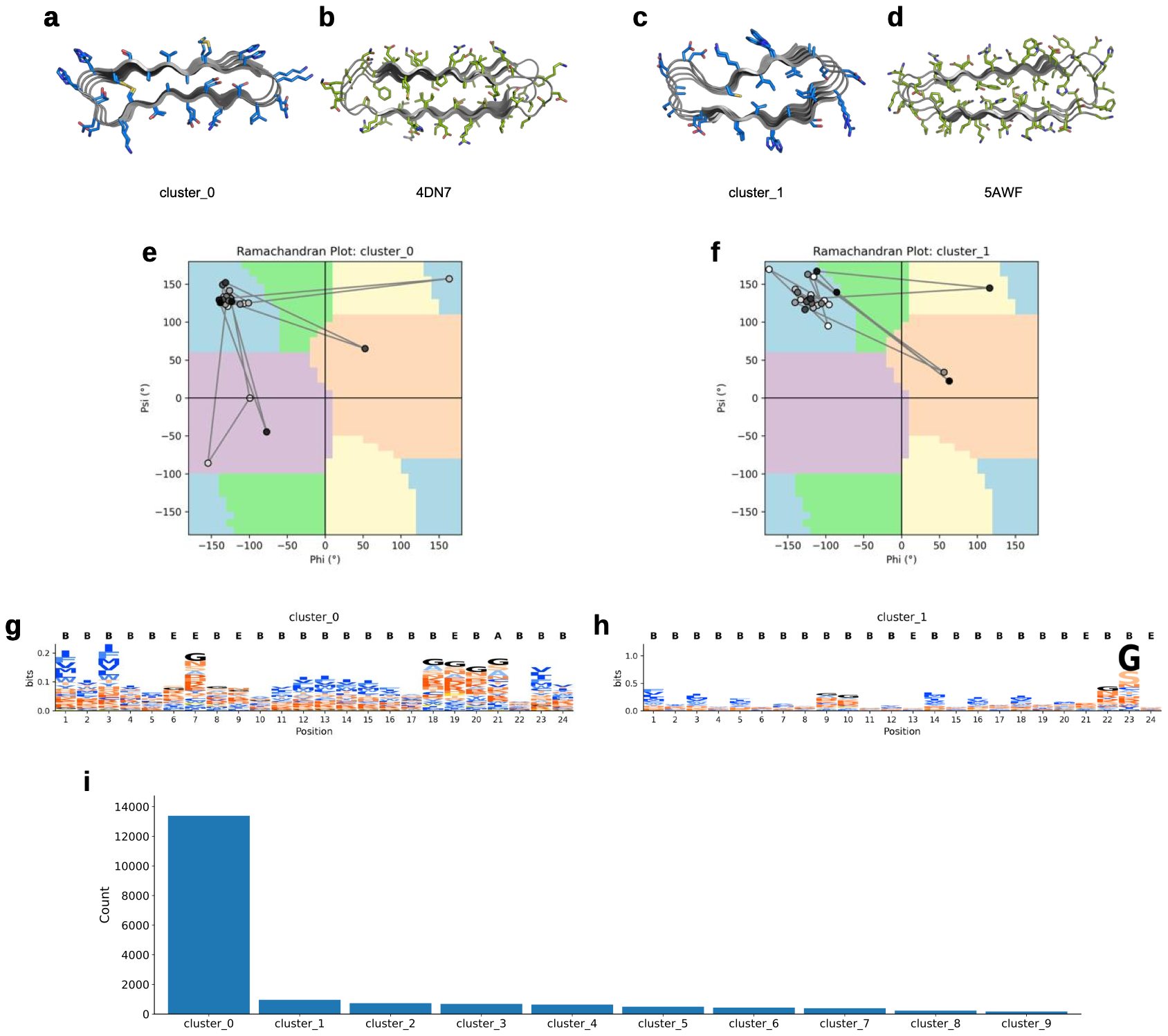

**Fig. S20. Conformational clustering and sequence features of designed 24-mer repeat proteins generated from the full amino acid set. (**A–D) Representative structures from cluster_0 and cluster_1 compared with their nearest PDB analogues (4DN7, 5AWF). (E, F) Ramachandran plots showing backbone angle distributions for clusters 0 and 1. (G, H) Sequence logos for each cluster, illustrating register-specific residue preferences across the 24-mer repeat. (I) Cluster size distribution for all designs.

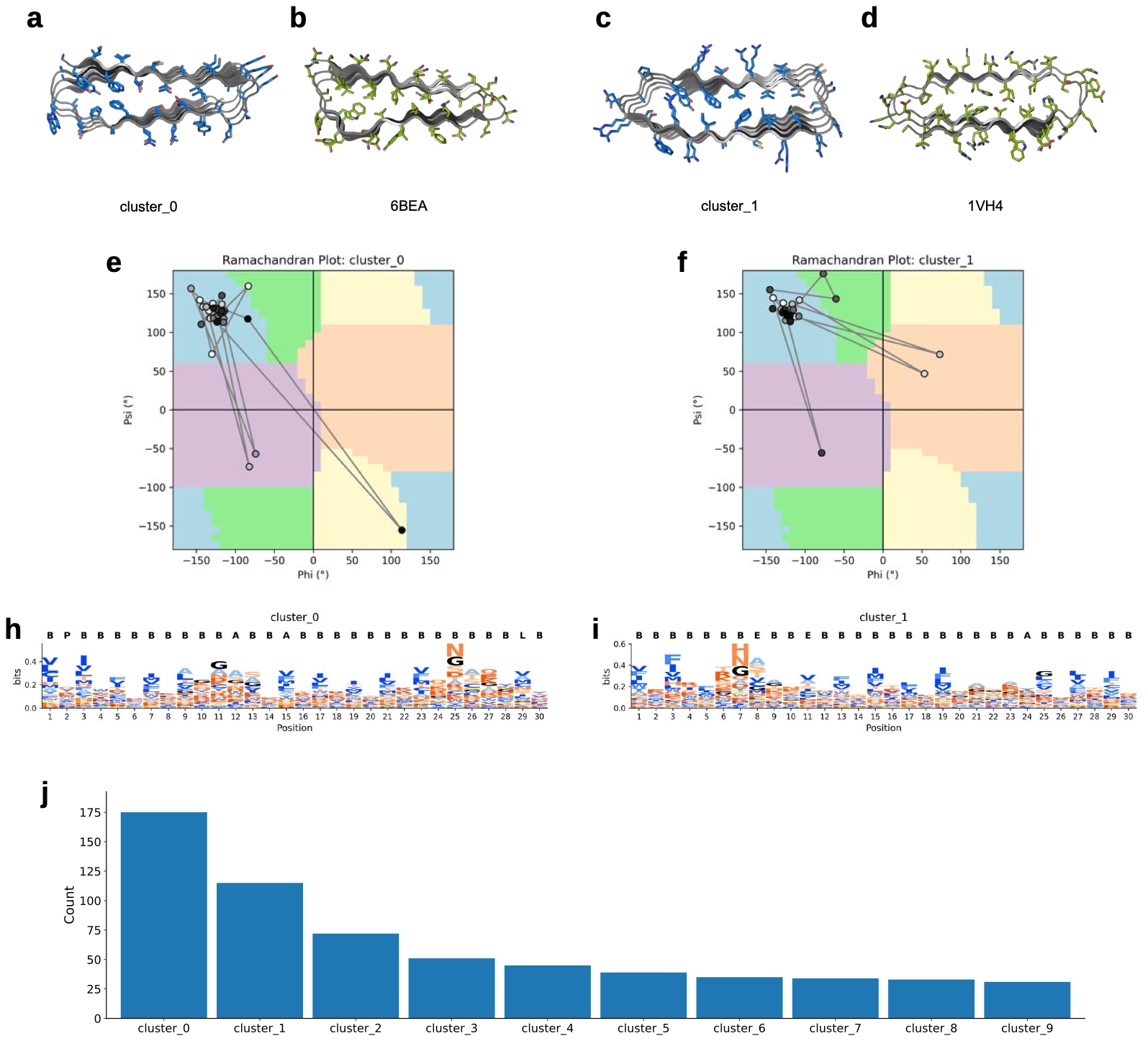

**Fig. S21. Conformational clustering and sequence features of designed 30-mer repeat proteins generated from the full amino acid set. (**A–D) Representative structures from cluster_0 and cluster_1 compared with their nearest PDB analogues (6BEA, 1VH4). (E, F) Ramachandran plots showing backbone angle distributions for clusters 0 and 1. (G, H) Sequence logos for each cluster, illustrating register-specific residue preferences across the 30-mer repeat. (I) Cluster size distribution for all designs.

>Heliscrew1

EEFLRLSPDEIAKLSPEEIAALTPEQIARLSPDKVAALTPGQIAALSPEQIAALSPEKVAAMTPDQVAALTPGQIAALSPEKIAALSPDVLRALTPEQWAALSPEQKAAFPADVRAELAR

>Heliscrew2

EEFLKLSPEEIRKLSPEEVAALTPDQIAALSPEKIRALTPEQLRALSPDQVAALSPEKIAALTPDQVRALTPEQIAALSPEKIAALSPDVLRALTPEQWAALTPEQKAAFPADVRAELAG

> Heliscrew3

EEFLAMTPEELAALSPEQIAALPVDVIARLSPDKIAGLTPEQIAALSPEQIAALSVDKIAALTPDQVRALTPEQIAALSPEKIAALSPDVIRALTPEQWAALSPEQKAALPADIRAELEG

>Heliscrew4

AEVAALTPEQIAALSVDEIARLSPDQFAALTPEQIAALTPEQVARLSPEKIAALSPDKIAAMTPEQVAALTPEQFAALSPDKIAALTPEQIAAVSDETLRALSPEKQAALPPEVRARLAG

>Heliscrew5

EEVAALTPEQVAALSPEKIAALSPEDFARLTPEQVAALTPEQIRALSPEKFAALSPEKIAALTPDQVAAMTPEQFAALSPEKIAALSPEQIAAVDMETLRALSPEKQAALPPEVRARLAG

>Heliscrew6

AMTPEEIAALSPEELRKLSPEQVRAWTPEQIAALSPDKIAALSPEQVRALTPEQIAALTPDKIAALSPEQVAGLTPEQIRALSPEKIAAISPETIARWPADVLAGLSPEQKAALPAALRA

>Heliscrew7 **(original cluster centroid)**

ALTVDQIKALTVDQIKALTVDQIKALTVDQIKALTVDQIKALTVDQIKALTVDQIKALTVDQIKALTVDQIKALTVDQIKALTVDQIKALTVDQIKALTVDQIKALTVDQIKALTVDQIK

**Fig. S22. Amino acid sequences of HeliScrews (1–7). HeliScrew7 corresponds to the original cluster centroid helical screw design.**

**>Heliscrew1**

**GAAGAATTCTTGCGCCTGAGCCCTGATGAGATCGCAAAACTGTCCCCGGAAGAAATAGCTGCCCTGACGCCCGAACAGATTGCGCGTCTTTCACCGGATAAAGTCGCCGCACTTACACCAGGCCAGATCGCTGCACTGAGTCCGGAGCAAATTGCGGCCCTCAGCCCAGAAAAAGTGGCCGCAATGACTCCGGACCAGGTGGCGGCGTTAACCCCGGGTCAGATTGCGGCCTTGAGCCCGGAAAAAATTGCGGCGTTATCTCCGGACGTACTGCGGGCGCTGACCCCCGAGCAGTGGGCTGCACTGTCGCCGGAACAAAAGGCGGCCTTTCCTGCCGATGTTCGCGCCGAGCTAGCGCGTTGATAA**

**>Heliscrew2**

**GAAGAATTTCTGAAACTGAGCCCCGAAGAGATCCGCAAACTGAGCCCCGAGGAGGTGGCAGCACTGACCCCGGACCAAATTGCAGCGCTTTCACCTGAAAAAATTCGTGCCCTGACACCGGAACAGCTGCGCGCGTTATCGCCTGATCAGGTCGCGGCGCTAAGTCCGGAAAAGATTGCTGCCCTGACCCCAGATCAAGTACGTGCATTAACCCCGGAGCAGATCGCGGCCCTTTCCCCGGAAAAGATTGCCGCTCTGTCTCCGGATGTGTTGCGAGCCCTCACGCCGGAACAGTGGGCGGCCCTCACTCCGGAACAGAAAGCGGCCTTCCCAGCGGACGTTCGCGCAGAATTGGCGGGCTGATAA**

**>Heliscrew3**

**GAGGAGTTTCTAGCCATGACACCGGAGGAATTGGCCGCACTCTCTCCCGAGCAGATTGCTGCGCTGCCTGTCGATGTGATTGCGCGTCTGTCCCCGGATAAAATTGCGGGCCTGACGCCAGAACAGATCGCCGCACTTTCACCGGAACAAATCGCGGCCCTGAGCGTGGACAAAATTGCGGCGCTGACCCCAGATCAGGTTCGCGCATTAACTCCTGAACAGATTGCGGCCTTGTCGCCGGAAAAGATTGCTGCCCTGAGTCCGGACGTAATCCGCGCCCTCACCCCCGAACAGTGGGCAGCATTAAGCCCGGAACAAAAAGCGGCTCTTCCGGCCGATATACGTGCGGAACTGGAAGGTTGATAA**

**>Heliscrew4**

**GCAGAAGTGGCAGCACTGACGCCGGAACAGATCGCGGCGTTGAGCGTGGACGAAATTGCCCGATTATCCCCAGATCAGTTTGCTGCCCTAACACCTGAACAGATCGCAGCGCTGACGCCGGAGCAAGTTGCGCGCCTGAGCCCTGAAAAAATTGCAGCCTTGTCTCCGGACAAGATTGCCGCAATGACCCCGGAGCAGGTTGCGGCCCTTACCCCAGAGCAGTTCGCGGCGCTGTCACCGGATAAAATAGCCGCGCTGACCCCCGAGCAAATTGCCGCTGTAAGTGATGAAACTTTACGCGCCCTTTCGCCCGAAAAACAGGCTGCCCTCCCGCCGGAAGTCCGTGCGCGTCTGGCGGGCTGATAA**

**>Heliscrew5**

**GAAGAAGTGGCAGCACTGACCCCTGAACAAGTGGCGGCGTTATCGCCGGAAAAAATTGCGGCCTTATCCCCCGAAGATTTTGCGCGCCTGACTCCTGAGCAGGTAGCGGCCCTCACGCCGGAACAGATCCGTGCCTTGAGCCCGGAAAAATTCGCGGCGCTGAGCCCGGAAAAAATCGCGGCTCTGACCCCGGATCAGGTTGCAGCCATGACACCGGAGCAGTTTGCGGCCTTGTCACCGGAGAAAATTGCAGCACTGAGTCCCGAACAGATTGCGGCCGTTGACATGGAAACCCTACGTGCTCTGTCTCCAGAGAAGCAAGCAGCTCTTCCGCCAGAGGTCCGAGCCCGCCTTGCCGGCTGATAA**

**>Heliscrew6**

**GCGCTGACGGTCGACCAGATTAAAGCACTGACTGTGGACCAGATTAAAGCTCTAACAGTTGATCAGATCAAGGCACTTACCGTCGATCAAATTAAAGCCCTCACGGTGGATCAAATTAAAGCCCTGACCGTCGATCAGATTAAGGCCTTGACCGTGGACCAAATCAAAGCGCTTACTGTAGATCAAATCAAGGCACTGACGGTTGATCAGATTAAAGCGTTAACCGTAGATCAGATTAAAGCGTTGACCGTGGACCAGATCAAAGCTCTGACGGTGGACCAGATAAAGGCGTTAACAGTTGATCAGATTAAAGCGCTGACCGTGGATCAGATTAAAGCCCTGACCGTTGATCAAATCAAATGATAA**

**>Heliscrew7 (original cluster centroid)**

**GCAATGACCCCGGAAGAGATTGCGGCATTGTCTCCAGAAGAACTGCGTAAACTTAGCCCGGAACAGGTCCGAGCGTGGACCCCGGAACAAATTGCGGCGCTGTCCCCTGATAAAATCGCCGCCCTCAGCCCGGAGCAGGTGCGCGCGTTAACACCGGAACAGATTGCGGCACTGACGCCAGATAAAATCGCGGCCCTAAGCCCCGAGCAGGTAGCCGGCCTGACTCCGGAACAGATACGTGCTTTATCGCCTGAGAAGATTGCAGCGATTTCACCGGAAACCATCGCTCGCTGGCCTGCCGACGTTCTTGCCGGTTTGAGTCCCGAACAAAAAGCGGCACTGCCGGCCGCCCTGCGCGCTTGATAA**

**Fig. S23. Codon optimized DNA sequences encoding HeliScrew1–7 used for recombinant expression in *E. coli*. HeliScrew7 corresponds to the original cluster centroid sequence.**

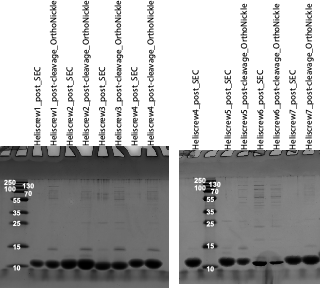

**Fig. S24. SDS-PAGE analysis of purified HeliScrew1–7. All seven designs express in *E. coli* and yield soluble proteins at the expected molecular weight. HeliScrew7 corresponds to the original cluster centroid design.**

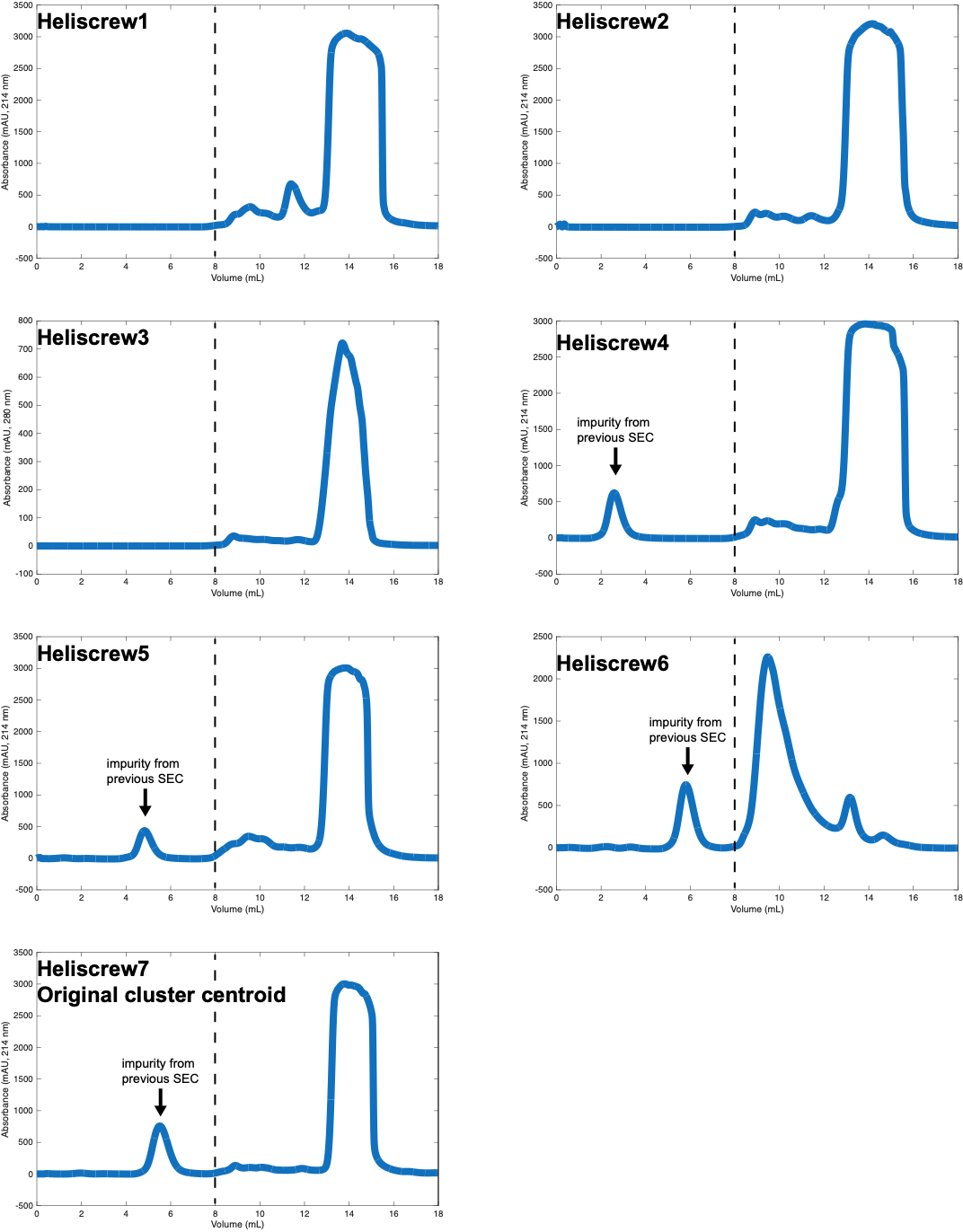

**Fig. S25. Size-exclusion chromatography (SEC) traces of HeliScrew1–7. Most designs elute as a predominant monomeric peak, whereas HeliScrew6 shows a higher molecular weight peak indicating higher-order assembly. HeliScrew7 corresponds to the original cluster centroid design.**

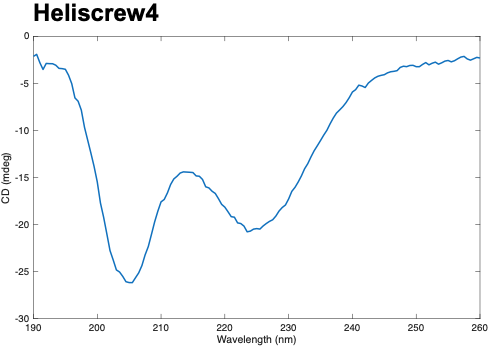

**Fig. S26. Far-UV circular dichroism spectrum of HeliScrew4. CD trace of HeliScrew4 diluted in H_2_O, showing a characteristic α-helical spectrum with negative minima at 208 and 222 nm.**

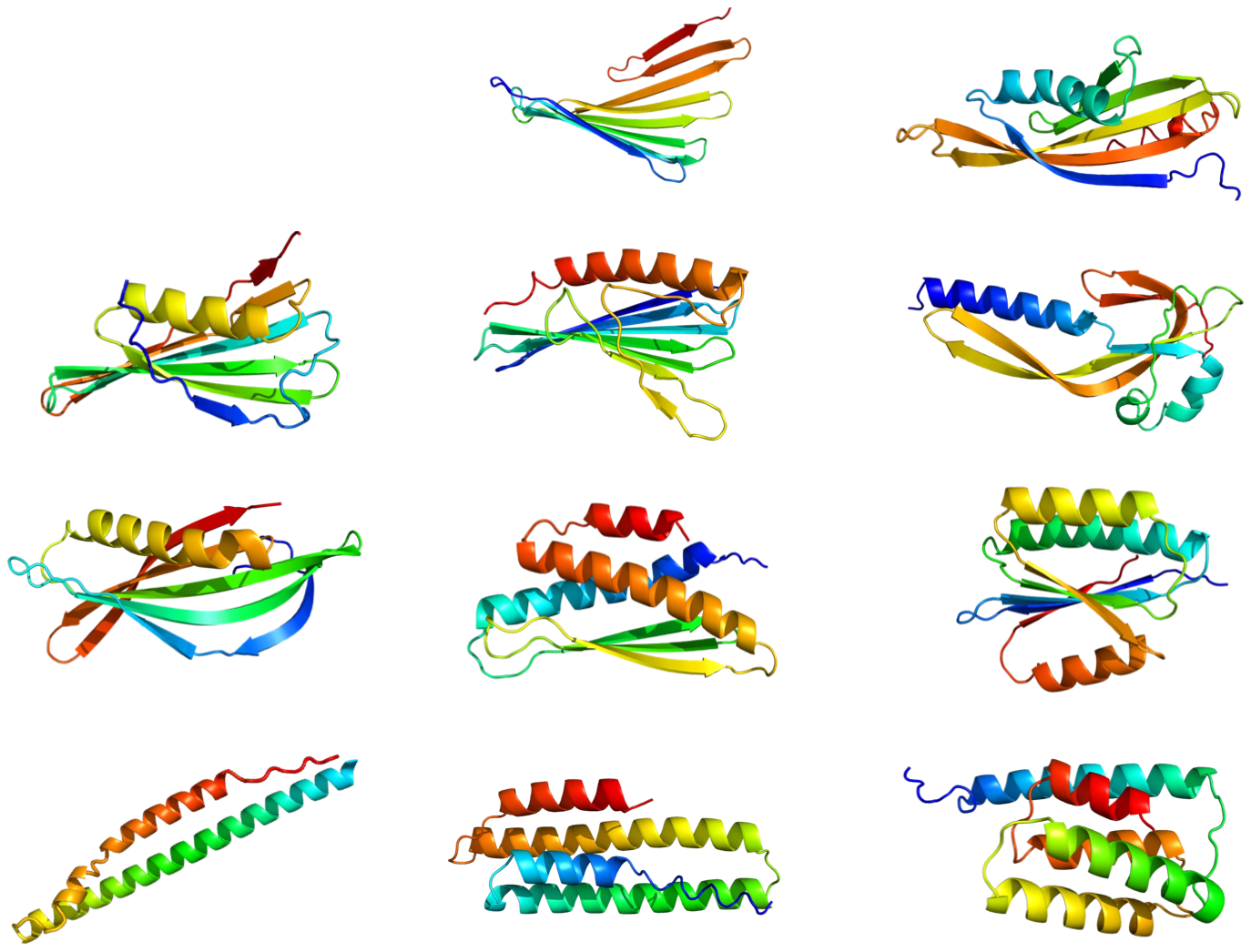

**Fig. S27.** Eleven designed 120-mer proteins predicted with pLDDT > 90.

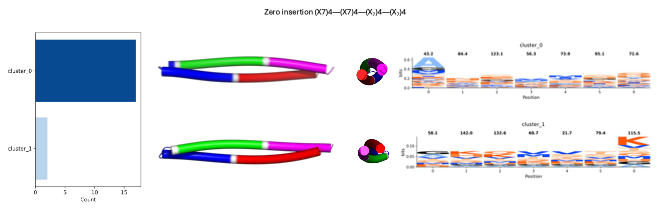

**Fig. S28. Conformational clustering and sequence features for the zero-insertion design.** Cluster populations (left), representative centroid structures (right) are shown for the two dominant clusters.

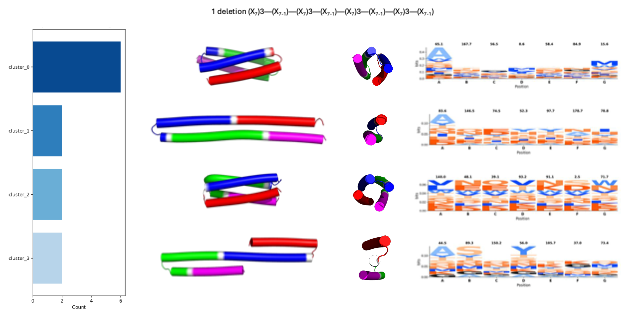

**Fig. S29. Conformational clustering and sequence features for 1-deletion.** Cluster populations (left), representative centroid structures (right) are shown for the four dominant clusters.

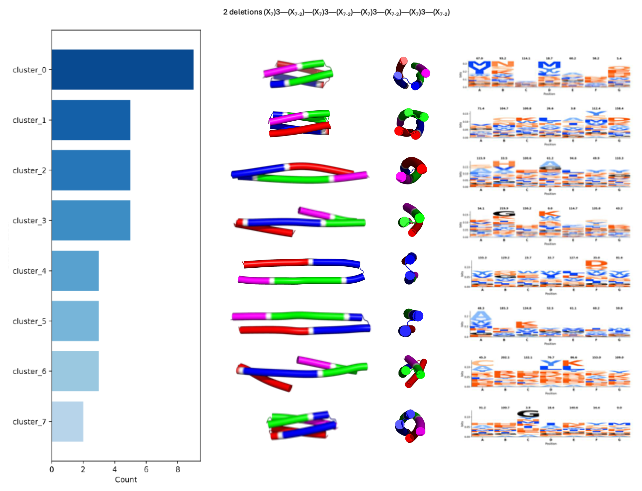

**Fig. S30. Conformational clustering and sequence features for 2-deletions.** Cluster populations (left), representative centroid structures (right) are shown for the eight dominant clusters.

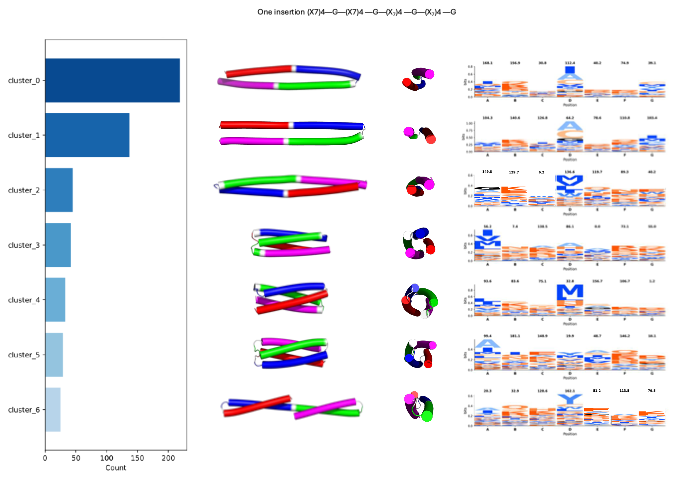

**Fig. S31. Conformational clustering and sequence features for 1-insertion.** Cluster populations (left), representative centroid structures (right) are shown for the seven dominant clusters.

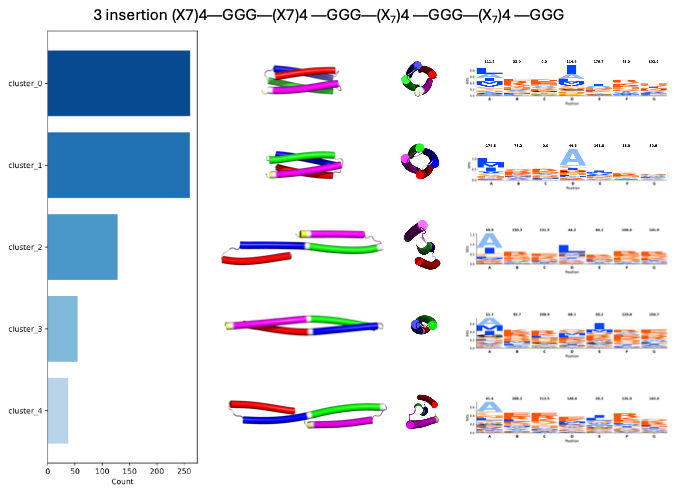

**Fig. S32. Conformational clustering and sequence features for 3-insertions.** Cluster populations (left), representative centroid structures (right) are shown for the five dominant clusters.

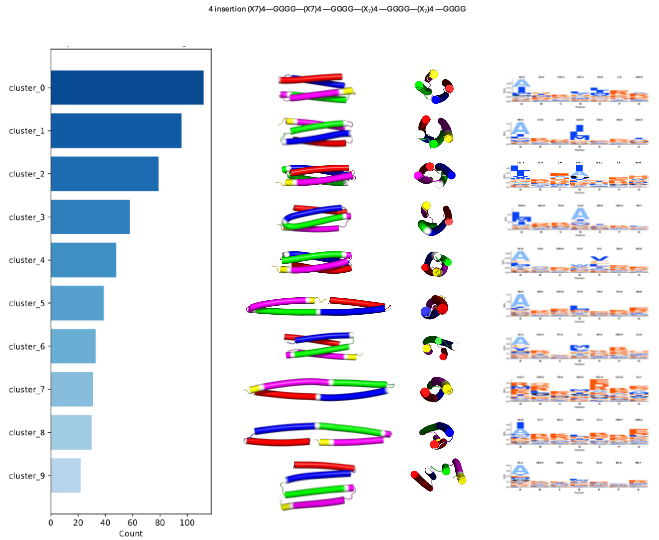

**Fig. S33. Conformational clustering and sequence features for 4-insertions.** Cluster populations (left), representative centroid structures (right) are shown for the ten dominant clusters.

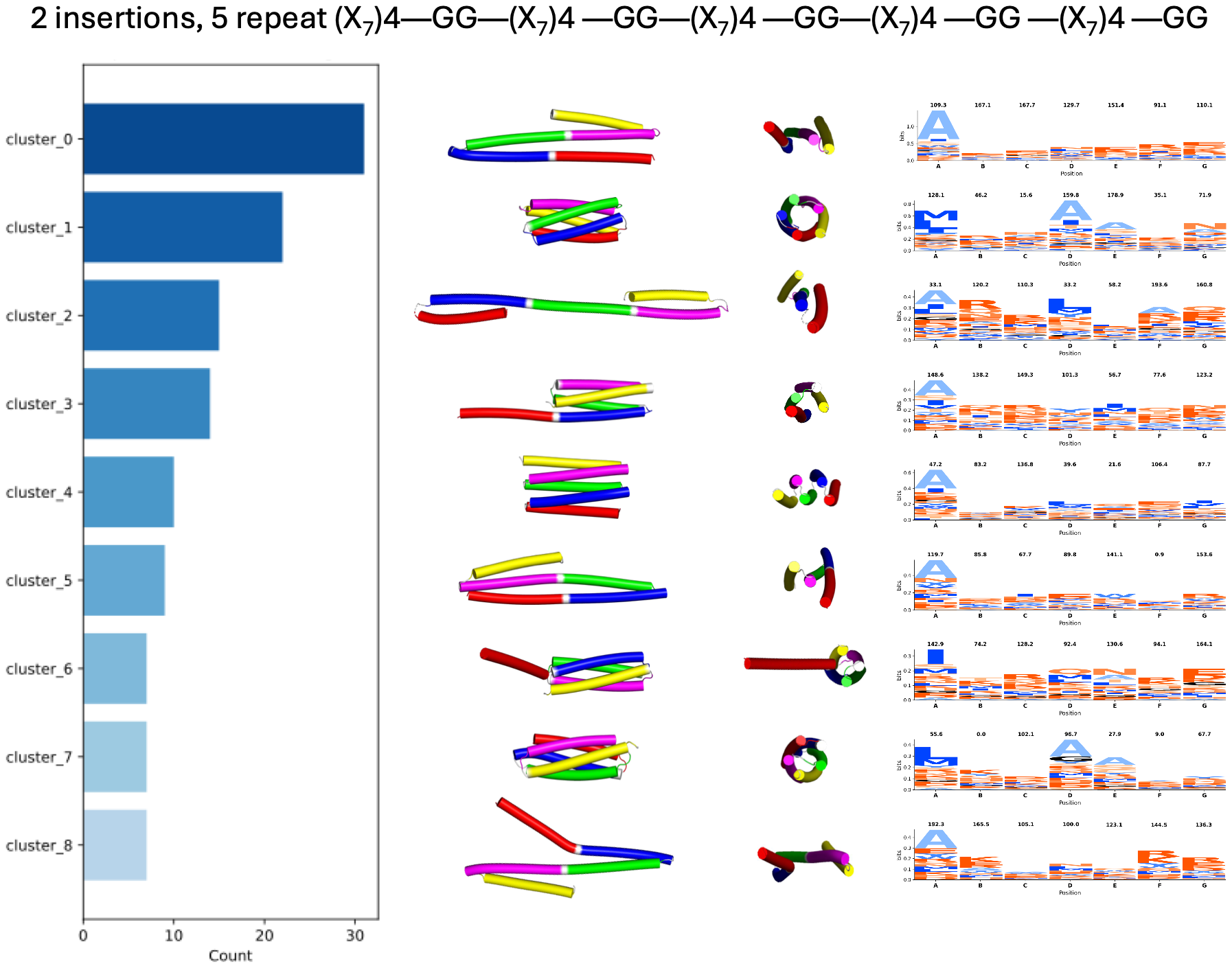

**Fig. S34. Conformational clustering and sequence features for 2-insertions, 5 repeats.** Cluster populations (left), representative centroid structures (right) are shown for the nine dominant clusters.

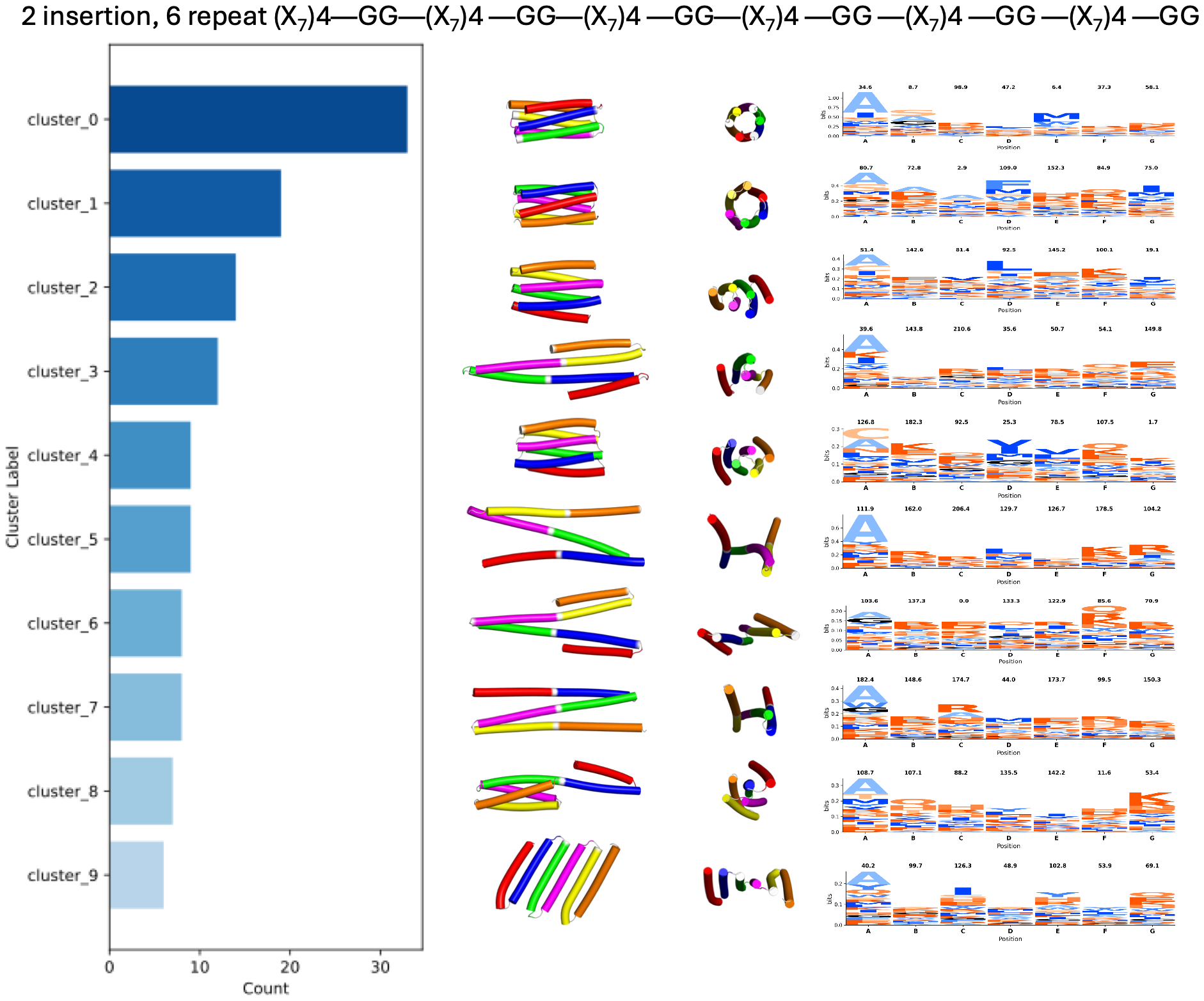

**Fig. S35. Conformational clustering and sequence features for 2-insertions, 6 repeats.** Cluster populations (left), representative centroid structures (right) are shown for the ten dominant clusters.

**Table S1. Top 10 cluster centroid structures across all repeat sizes.** Representative centroid structures for the ten most populated clusters are shown for each repeat length (5mer–40mer).

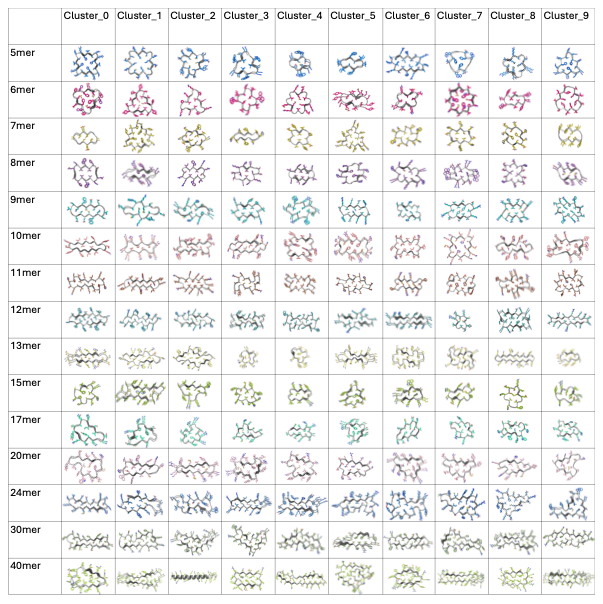

**Table S2.** Data collection and refinement statistics of Heliscrew4.

|  | **Heliscrew4* (PDB 9ZEM)** |
| --- | --- |
| **Wavelength** | 0.980 |
| **Resolution range** | 45.69 - 2.046 (2.09 - 2.05) |
| **Space group** | P 1 21 1 |
| **Unit cell** | 37.339 62.62 45.98 90 96.42 90 |
| **Total reflections** | 68979 (4248) |
| **Unique reflections** | 12754 (765) |
| **Multiplicity** | 5.4 (5.6) |
| **Completeness (%)** | 92.43 (91.24) |
| **Mean I/sigma(I)** | 5.29 (3.17) |
| **Wilson B-factor** | 18.02 |
| **R-merge** | 0.311 (1.113) |
| **R-meas** | 0.3437 (1.228) |
| **R-pim** | 0.1432 (0.5093) |
| **CC1/2** | 0.95 (0.392) |
| **CC*** | 0.987 (0.751) |
| **Reflections used in refinement** | 12731 (762) |
| **Reflections used for R-free** | 1272 (77) |
| **R-work** | 0.2088 (0.3069) |
| **R-free** | 0.2438 (0.3217) |
| **Number of non-hydrogen atoms** | 1869 |
| **macromolecules** | 1718 |
| **ligands** | 0 |
| **solvent** | 151 |
| **Protein residues** | 234 |
| **RMS(bonds)** | 0.006 |
| **RMS(angles)** | 0.91 |
| **Ramachandran favored (%)** | 100.00 |
| **Ramachandran allowed (%)** | 0.00 |
| **Ramachandran outliers (%)** | 0.00 |
| **Rotamer outliers (%)** | 0.00 |
| **Clashscore** | 6.86 |
| **Average B-factor** | 24.84 |
| **macromolecules** | 24.36 |
| **solvent** | 30.32 |

Statistics for the highest-resolution shell are shown in parentheses.
